# Recent ultra-rare inherited mutations identify novel autism candidate risk genes

**DOI:** 10.1101/2020.02.10.932327

**Authors:** Amy B Wilfert, Tychele N Turner, Shwetha C Murali, PingHsun Hsieh, Arvis Sulovari, Tianyun Wang, Bradley P Coe, Hui Guo, Kendra Hoekzema, Trygve E Bakken, Lara H Winterkorn, Uday S Evani, Marta Byrska-Bishop, Rachel K Earl, Raphael A Bernier, The SPARK Consortium, Michael C Zody, Evan E Eichler

## Abstract

Autism is a highly heritable, complex disorder where *de novo* mutation (DNM) variation contributes significantly to disease risk. Using whole-genome sequencing data from 3,474 families, we investigate another source of large-effect risk variation, ultra-rare mutations. We report and replicate a transmission disequilibrium of private likely-gene disruptive (LGD) mutations in probands but find that 95% of this burden resides outside of known DNM-enriched genes. This variant class more strongly affects multiplex family probands and supports a multi-hit model for autism. Candidate genes with private LGD variants preferentially transmitted to probands converge on the E3 ubiquitin-protein ligase complex, intracellular transport, and Erb signaling protein networks. We estimate these mutations are ~2.5 generations old and significantly younger than other mutations of similar type and frequency in siblings. Overall, private LGD variants are under strong purifying selection and act on a distinct set of genes not yet associated with autism.

**One sentence summary:** Ultra-rare autism variants preferentially transmitted to probands are younger and identify distinct gene candidates and functional networks.

---

Autism spectrum disorder (ASD) is a highly phenotypically heterogeneous disorder that affects about 1 in 59 children in the United States (*1*). Studies to date have primarily focused on high-impact and sporadic variants, such as *de novo* copy number variants (CNVs) and single-nucleotide variants (SNVs). Despite their large effect sizes, each category of *de novo* mutation (DNM) only accounts for a small fraction of cases with the upper bound of autism cases accounted for by DNM estimated to be ~25% (*2–4*). Although this genetic model is highly relevant to simplex ASD, where there is only one affected child in a family, it does not explain most cases and is less likely for most multiplex families, or families with more than one affected child (*5*). This has led to the reassessment of the role of various classes of inherited mutation and their contribution to autism risk (*3, 6–9*).

For example, it is well established that large and sometimes inherited CNVs contribute to a small percentage of both sporadic and multiplex autism (*3, 4*). Preferential transmission of likely gene-disruptive (LGD) mutations have been observed in both simplex (*3*) and multiplex autism families (*7*) whose genes have been reported to converge on genes in related functional networks (*3, 7*). Also, genetic studies of ASD and developmental delay families have found evidence of an enrichment of multiple gene-disruptive mutations (both CNVs and SNVs) in the affected children (*10–13*). In addition, recent analyses suggest that common inherited risk variants also contribute at some level to most ASD pathology (*14, 15*); however, it is unclear whether common risk variants alone are sufficient to cause disease.

In this study, we focus on ultra-rare mutations that are unique to a family and are likely to disrupt gene function. We expand the number of multiplex and simplex autism families sequenced taking advantage of the increased sensitivity afforded by whole-genome sequencing (WGS) over whole-exome sequencing (WES) data (*16*) to create a highly sensitive LGD variant callset from WGS data from 3,474 simplex and multiplex autism families from the Centers for Common Disease Genomics (CCDG) (Table 1 **and table S1**). We use these to assess transmission biases of this class of variant to probands and unaffected siblings after controlling for population structure (*7, 17–20*). Further, we replicate these analyses in WES data from 5,879 families from the Simons Foundation Powering Autism Research Knowledge (SPARK) (*21, 22*). Our results provide strong support for private LGD mutations contributing to autism and, in particular, multiple hits in different genes. We show that such mutations have arisen more recently in autism families (2.5 generations) when compared to other classes of variants. Importantly, genes associated with DNM enrichment contribute little to this burden, rather our analyses identify new gene candidates enriched in specific functional pathways.

**Table 1:** Summary of whole-genome and whole-exome sequencing

| **Cohort** | **Individuals** |  |  |  | **Families** |  |  | **Publication** | **Assay** |
| --- | --- | --- | --- | --- | --- | --- | --- | --- | --- |
|  | **Genomes** | **Probands** | **Siblings** | **Twin pairs** | **Simplex** | **Multiplex** | **Total** |  |  |
| **SSC** | 8757 | 2299 | 1860 | 1 | 2299 | 0 | 2299 | An et al. | WGS |
| **SAGE** | 547 | 202 | 5 | 4 | 144 | 26 | 170 | Guo et al. | WGS |
| **TASC** | 750 | 250 | 0 | 1 | 250 | 0 | 250 | Unpublished | WGS |
| **AGRE/**<br>**iHart Phase I\*** | 1736 | 822 | 194 | 69 | 6 | 354 | 360 | Ruzzo et al. | WGS |
| **AGRE/**<br>**iHart Phase II** | 1757 | 791 | 176 | 0 | 1 | 394 | 395 | Unpublished | WGS |
| **Discovery set** | 13547 | 4364 | 2235 | 75 | 2700 | 774 | 3474 | - | WGS |
| **SPARK** | 21331 | 6539 | 3034 | 63 | 5278 | 601 | 5879 | Unpublished | WES |
| **Combined** | 34880 | 10905 | 5269 | 164 | 7978 | 1375 | 9353 | - | WGS & WES |

Summary information of the five cohorts included in our study. WGS samples have been sequenced to an average depth of 34x.
\*10 families had additional members sequenced in AGRE/iHart Phase II. These families were still included in the number of simplex/multiplex families for AGRE/iHart phase I; however, the family was identified as simplex or multiplex based on the full family data, which includes the additional members from AGRE/iHart Phase II.

## Whole-genome sequencing

We generated WGS data (30-fold sequence coverage) from 2,507 individual DNA samples from 394 multiplex and 251 simplex autism families (Table 1, **table S1, and Materials and Methods**). Combined with previously published WGS data (*7, 18, 19*), we created a standardized callset of SNVs and indels from 13,547 genome samples using two callers and have made these publically available (**Data and materials availability**). The set consists of data from 4,364 probands and 2,235 siblings and includes parent–child SNV data from 774 multiplex and 2,700 simplex families. Focusing initially on DNMs, we performed 582 random Sanger sequencing validation experiments and combined these with previously published validation experiments (**Table S2**) (*4*). We report an overall validation rate of 99.5% for our DNM callset and estimate a false negative rate of 3.5%. On average, we observe 65.14 DNMs per child and an increase of 1.11 and 0.37 mutations per year of paternal and maternal age, respectively (**Fig. S1**).

## Discovery and properties of private variants

While previous studies have focused on the contribution of DNMs or common variants underlying ASD (*2–5*), we sought to reevaluate the contribution of transmitted variants to the genetic etiology of ASD (*3, 7*). Because our previous study showed that transmission disequilibrium signals increased with rarer inherited variants (*3*), we specifically focused on private inherited variants. We define these as heterozygous variants that are observed only once in the parent population and transmitted to at least one child, regardless of potential *de novo* status in unrelated children within the cohort. We note, for example, that 0.036% of our private mutations overlap with mutations in our DNM callset. Based on our sample size, private variants correspond to an approximate allele frequency ≤ 7e^−5^ and are ultra-rare in nature. We identified 26,606,722 unique private variants (35,871,117 total) in our discovery cohort of 6,599 children and detected no difference in the average number of private variants between probands and unaffected siblings genome-wide (Mann-Whitney test, autosomes: p = 0.168, female X chromosome: p = 0.328, male X chromosome: p = 0.534). We detect no difference in the number of private autosomal variants transmitted from fathers as compared to mothers genome-wide (Wilcoxon signed-rank test, p = 0.1995) but did detect a difference, as expected, if we consider the female X chromosome: Wilcoxon signed-rank test, p = 0.0215; mean paternally transmitted: 98.7, mean maternally transmitted: 102).

Since individuals of similar ancestry will have an increased chance of allele sharing as compared to individuals from different populations (*23*), we considered private mutations in the context of ancestry. We assigned each individual to one of six super populations (EUR, AFR, AMR, EAS, SAS, and OCN) based on maximum likelihood estimations of ancestry using a human diversity panel (**Materials and Methods, Fig. S2A**). Consistent with previous studies (*24, 25*), children of European descent carry the fewest private variants per genome (Fig. 1, B and C **and tables S3 and S4**). This is because the majority of subjects in the discovery cohort (85.6%) are of European descent (**Fig. S2B**). Thus, private variants among the EUR subgroup will be of the lowest frequency providing the greatest specificity, in principle, to detect pathogenic events in this study (*3*).

**Fig. 1:**
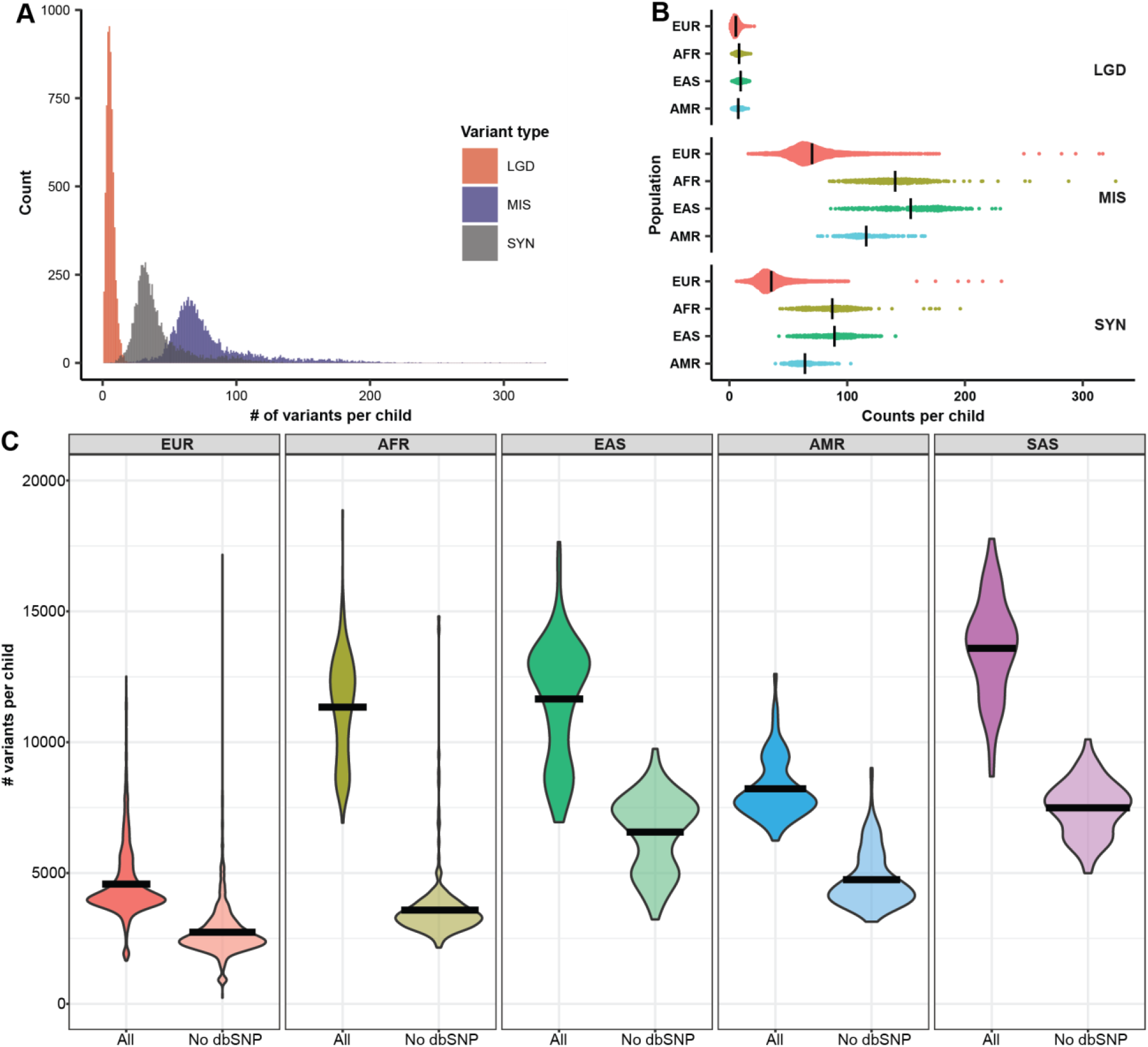
Overview of private variants in discovery cohort. Private variants are defined as variants observed in one and only one parent in the cohort. (A) Distribution of likely gene-disruptive (LGD), missense (MIS) and synonymous (SYN) private mutations per child (probands and unaffected siblings). (B) The cumulative number of each variant class by assigned population group (EUR=European (n = 5,685), AFR=African (n = 290), EAS=East Asian descent (n = 252), AMR=Amerindian (n = 193), SAS=South Asian (n = 103)), excluding SAS. (C) Private, transmitted variant counts per child grouped by ancestry before (All) and after filtering with dbSNPv150 (No dbSNP). Excess of private variants is partially but not fully resolved after excluding sites observed in dbSNP. We were unable to assign ancestry to one of these five population groups for 74 of the children in this study; y-axis was truncated at 20,000 variants per child; however, both the AFR and EUR populations had a small number of children with variant counts above this threshold (see tables S2 and S3 for details).

We tested whether filtering against a genetic database (dbSNP150) would be sufficient to eliminate this effect and improve our specificity for private events in the other populations (Fig. 1C, **fig. S3, and table S4**). Although dbSNP filtering did reduce the average number of private variants per child across populations, the magnitude of the effect varied by population. For example, among Africans, where diversity is the greatest, this treatment reduced by 69.2% the number of private variants but led to a reduction of only 43.6% and 44.9% among East and South Asian populations, respectively. We note that African and East Asian children had, on average, similar variant counts prior to dbSNP filtering (mean: 11,630 AFR vs. 11,653 EAS, **Table S4**). This suggests that the composition of the population genetic database may, in fact, introduce additional biases for populations because sampling has been nonuniform and that allele frequency filtering alone is not sufficient to account for population stratification. These differences highlight the need to account for ancestry and to increase representation of underrepresented populations for gene discovery—even for rare variant analyses. Therefore, all analyses reported in this study have been replicated in the SPARK cohort and confirmed in the European subset of our discovery cohort.

## Patterns of private, transmitted variants in protein-coding regions

In this study, we restrict our analyses to autosomal, protein-coding regions of the genome, where we expect to have the greatest power to detect enrichment of private, transmitted variants (*2, 3, 7*). Missense mutations are the most abundant followed by synonymous and then LGD mutations, defined here as stop-gain, stop-loss, splice-altering SNV or frameshift indels (Fig. 1A **and table S4**). We observe no significant difference between the overall proportion of proband and unaffected sibling carriers for any of the three mutation classes and detect no significant enrichment when considering all genes (LGD: Fisher’s exact test OR = 1.08, p = 0.792, MIS: OR = 0, p = 1,SYN: OR = 0 p = 1). However, when we consider subsets of genes at increasing thresholds of gene constraint using pLI, we replicate (*3, 7, 26, 27*) the relationship of increasing transmission of LGD mutations to probands with increasing gene constraint for the discovery, replication, and combined cohort (Fig. 2A, **fig. S4, and table S5**). This is not observed for missense or synonymous mutations. Interestingly, if we partition our discovery cohort by sex to include the X chromosome, we find that the increased burden of private LGD mutations relative to increasing gene constraint becomes both stronger and more significant in males (**Figs. S5 and S6A and table S5**, pLI ≥ 0.90, male vs. female OR = 1.36 vs. 1.10, permutation p-value = 0.004).

**Fig. 2:**
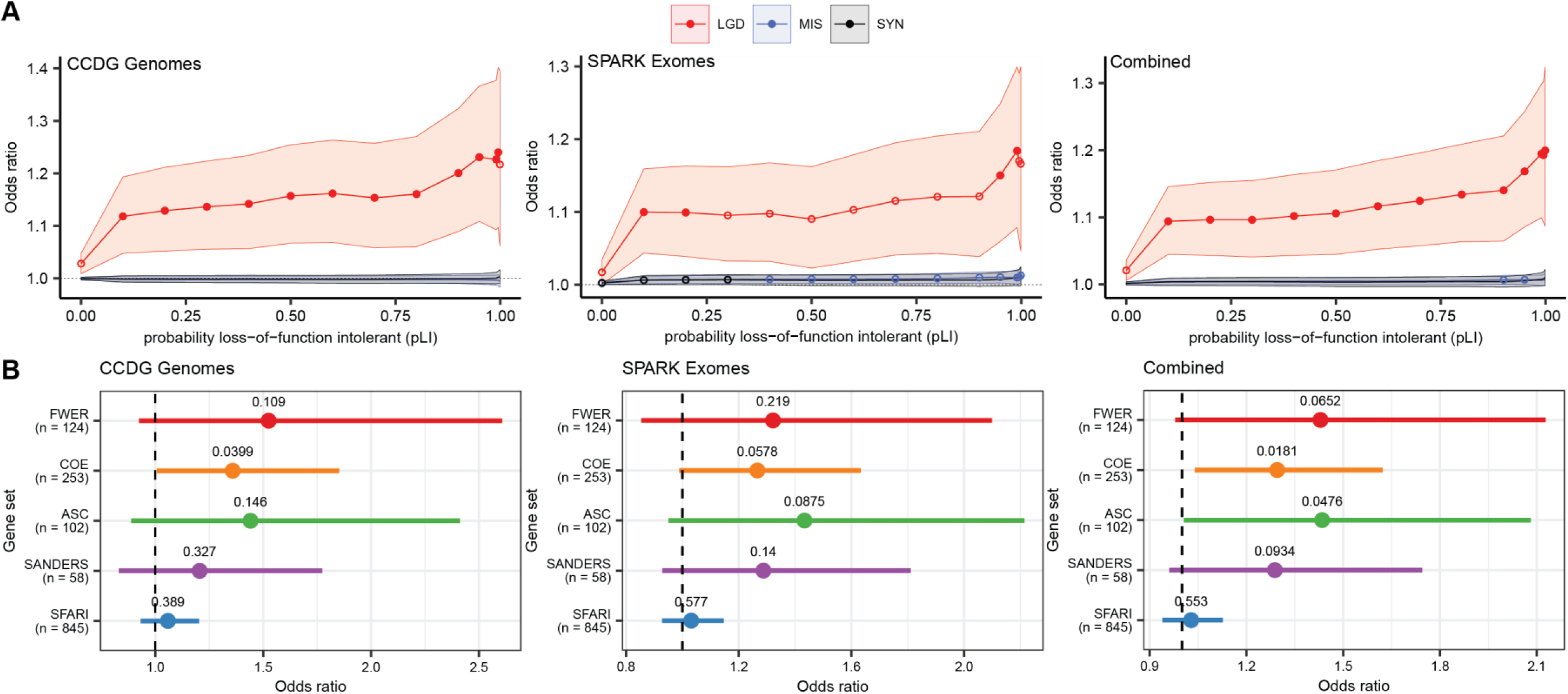
Burden of private LGD variants in affected children. (A) Burden of private LGD mutations in probands as compared to siblings was quantified (odds ratio [OR]) at increasing thresholds of gene constraint (pLI) in our discovery (n = 4,201 affected and 2,191 unaffected children), replication (n = 6,453 affected and 3,007 unaffected children), and combined discovery and replication (n = 10,657 affected and 5,199 unaffected children) cohorts. Filled circles indicate Bonferroni-corrected p-values < 0.05 (42 tests per cohort). Unfilled circles indicated nominal p-values < 0.05. OR confidence intervals were calculated using logistic regression (see table S4 for details). (B) Enrichment of private LGD variant transmission to probands for five autism risk gene sets (FWER, COE, ASC, SANDERS, SFARI). With the exception of SFARI, most gene sets were identified based on an excess of *de novo* mutations (DNMs) in parent–child trios (see Materials and Methods). OR was based on a comparison of the proportion of carriers between probands and siblings in our discovery (n = 4,201 affected and 2,191 unaffected children), replication (n = 6,453 affected and 3,007 unaffected children), and combined (n = 10,657 affected and 5,199 unaffected children) cohorts using a Fisher’s exact test (see table S5 for details). Dashed black line indicates OR = 1, which represents no difference between probands and siblings. Families with monozygotic twins (n = 75 in discovery, n = 63 in replication, and n = 138 in combined) were removed from analysis. For the combined set, variants were restricted to regions with at least 20x average coverage in the exomes. Reported p-values are nominal.

Next, we compared transmission disequilibrium of private LGD variants among various autism risk gene sets. These included genes shown to be enriched for DNMs in ASD and neurodevelopmental disorder (NDD) cases (*9, 28, 29*) and 845 genes from Simons Foundation Autism Research Initiative (SFARI) (**Methods and Materials**). All genes show a trend toward enrichment of private LGD variants among probands when compared to unaffected siblings but to varying degrees. For example, the Coe gene set (*28*) shows nominal significance for enrichment in our discovery (Fig. 2B, **fig. S7, and tables S6 and S7**, OR = 1.36, nominal p = 0.040) and combined (Fig. 2B **and tables S6 and S7**, OR = 1.29, nominal p = 0.018) cohorts. In fact, it also reaches nominal significance in females (**Fig. S8 and tables S6 and S7**, OR = 1.71, nominal p = 0.0316) if we assess the burden in males and females separately to include genes on the X chromosome. Importantly, the trends are consistent between replication and discovery cohorts suggesting that only larger sample sizes are required to achieve significance that survives multiple-test correction. In general, DNM-derived gene sets show greater enrichment than a more general set of autism risk genes (i.e., SFARI). DNM-enriched gene sets derived from ASD and NDD studies perform as well (if not better) than those derived strictly from autism cohorts. Importantly, all trends disappear if we consider variants at higher allele counts in the parent population (**Fig. S9**), indicating that the signal is specific to ultra-rare mutations.

Based on the initial sequencing of the SPARK autism families, Feliciano et al. reported that most of the rare LGD mutation transmission bias could not be accounted for by known ASD/NDD genes (*21*). We reevaluated the burden of private, transmitted LGD mutations at increasing thresholds of gene constraint excluding genes enriched for DNMs in order to better quantify this effect. We find that 95.4% of private, transmitted LGD mutation burden in probands remains (Fig. 3A **and table S8**) at a pLI ≥ 0.99. Moreover, we estimate that private LGD variants in these DNM-enriched genes account for 1.45% of ASD risk, whereas private, transmitted LGD mutations in the remaining genes at a pLI ≥ 0.99 account for 2.64% of ASD risk (Table 2). Unlike putative truncating mutations associated with DNM, we estimate that most of the attributable risk for private mutations awaits discovery and these will be identified among genes not already associated with DNM burden. Taken together, these results confirm that the DNM-enriched genes are highly specific to ASD and confer substantial risk; however, there is additional burden in the less specific but more sensitive set of constrained genes (pLI ≥ 0.99) that is yet to be discovered.

**Table 2:** Population attributable risk for *de novo* and private LGD mutations

| **Mutation class** | **ASD genes**<br>**(DNM enriched)** | **Remaining genes,**<br>**pLI ≥ 0.99** |
| --- | --- | --- |
| *De novo* * | 4.39% | 1.45% |
| Private | 1.45% | 2.64% |

Population attributable risk (PAR) percentages were calculated in our discovery cohort for *de novo* and private LGD mutations in children (Methods). DNM calculations do not include the AGRE study. We defined the DNM-enriched ASD gene set as the genes reported in Coe et al., Sanders et al., and Satterstrom et al. \*Does not include AGRE cohort.

**Fig. 3:**
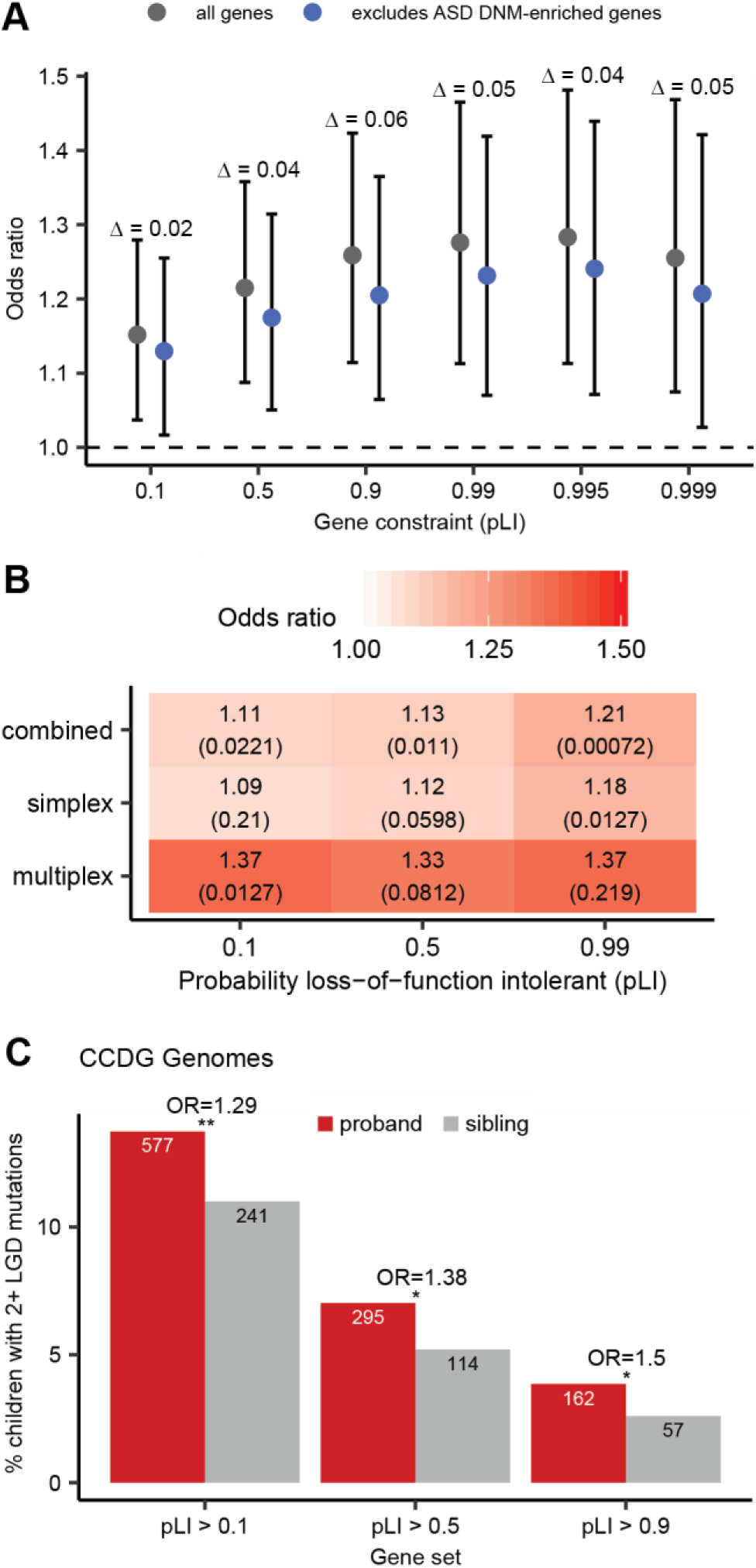
Genetic properties of inherited LGD mutation burden. (A) At least 95.4% of private, transmitted LGD mutation burden resides outside of autism risk genes identified with an excess of DNMs (321 genes considered and 154 genes with transmissions) based on analysis of CCDG autism genomes (n = 4,201 affected and 2,191 unaffected children). We observe 141 DNM-enriched genes with transmissions to probands and 85 genes with transmissions to siblings (Table S12). OR for five cumulative pLI bins were compared before and after excluding DNM-enriched ASD genes. The percentage of remaining burden is calculated as quotient of the OR for the pLI bin after removing DNM-enriched ASD genes and the OR for that pLI bin prior to removing the DNM-enriched ASD genes. Families with monozygotic twins (n = 75) were excluded from this analysis. OR and associated p-values were calculated using a Fisher’s exact test. (B) Multiplex families (n = 1,268 families, 2,691 probands, 533 siblings) show a higher burden of private, transmitted LGD mutations in probands as compared to siblings across three pLI thresholds compared to simplex families (n = 7,962 families, 7,962 probands, 4,666 siblings). (C) We observe a significant enrichment of probands carrying two or more private, transmitted LGD mutations (2+ LGD) when compared to unaffected siblings at various levels of gene constraint (3 cumulative pLI bins considered) based on CCDG genomes sequenced from autism families (n = 4,201 probands, 2,191 siblings). Families with monozygotic twins (n = 75) were excluded from this analysis. OR was calculated using Fisher’s exact test and reported p-values are Bonferroni corrected for (B) nine and (C) three tests (see Tables S7, S8 for details).

## Simplex versus multiplex and a multi-hit model for ASD

Both discovery and replication cohorts consist of a mixture of simplex families (only one affected child) and multiplex families (two or more affected children). Simplex families are hypothesized and have been shown (*2, 30*) to be enriched for sporadic or *de novo* genetic events (*17*), while multiplex families are more likely to inherit disease-causing mutations(*31*). We compared the proportion of proband versus siblings carrying at least one private LGD variant at increasing thresholds of gene constraint considering simplex and multiplex families independently (n = 2,700 simplex vs. 774 multiplex families, Table 1 **and table S1**). Despite having 3.5-fold fewer families, multiplex families show a 25.7% higher burden of private, transmitted LGD variants in probands when compared to simplex families with the greatest effect in multiplex families observed when considering unconstrained genes (Fig. 3B, **figs. S6B and S10, and table S9**, multiplex vs. simplex OR = 1.37 vs. 1.09, permuted p-value = 0.004 at a pLI ≥ 0.1). Among simplex families, significant burden is observed, in contrast, only among genes intolerant to mutation (pLI > 0.99).

Previous CNV work and analysis of putative noncoding DNMs (*11, 16*) have shown that autism probands are enriched for multiple deleterious mutations. If the signal we observed was relevant to the genetic etiology of autism, we hypothesized that affected children would be more likely to carry multiple private LGD mutations, partially explaining why both parents are unaffected in simplex families or are less severely affected in multiplex families. To this end, we compared the transmission of two or more private LGD mutations in probands and unaffected siblings conditioning on intolerance to mutation. We find that probands are significantly more likely to carry multiple inherited, LGD mutations in relatively unconstrained genes when compared to unaffected siblings (Fig. 3C, **fig. S11, and table S10**, OR = 1.29, Bonferroni-corrected p = 0.026 at pLI ≥ 0.1). Under an additive model, which represents independent assortment and random segregation, we would expect the odds ratio for the two-hit model to equal the square of the odds ratio for the one-hit model. Indeed, this is exactly what we observe. Additionally, this effect becomes stronger if we restrict the analysis to individuals of European descent (**Fig. S12**), indicating that this signal is not an artifact of population stratification.

## Novel candidate genes and interconnected functional networks

We investigated whether highly constrained genes that have not been reported as enriched for DNMs showed any enrichment for expression or protein-protein interaction (PPI) networks. Previous studies (*7, 9, 32*) have typically performed such analyses by integrating such candidates with DNM-enriched genes as opposed to considering them separately. Here, we focus on 163 highly constrained genes (**Table S11**) where private LGD mutations are exclusively transmitted to probands and have not been previously reported in SFARI or as DNM enriched in three ASD/NDD studies (*9, 28, 29*). Among the 163 genes, there are a total of 276 LGD mutations and 28 genes that have two or more independent LGD mutations observed in unrelated families.

Gene ontology analysis shows that the candidate gene set is highly enriched for encoded phosphoproteins (**Table S12**, KW-0597, 129/163 genes, q = 1.93e-20) and that the genes are more likely to be interconnected as part of PPI networks (Fig. 4, 102 observed vs. 75 expected edges, p-value = 0.00164). A subset of the genes (74/163 genes), including half of those with independent events in multiple families, converge on several functional pathways (Fig. 4 **and table S12**). This includes a small network of genes enriched for the E3 ubiquitin-protein ligase pathway by both the GO and Reactome databases, which are involved in proteasome degradation (HSA-98316) and regulation of protein modification by small protein conjugation or removal (GO:1903320). Similarly, there is a set of more than dozen genes associated with internal cellular transport and, in particular, transport between the Golgi and endoplasmic reticulum. Other subnetworks are significantly enriched for nucleobase-containing compound metabolic process (GO:006139) and Erb signaling (hsa04012).

**Fig. 4:**
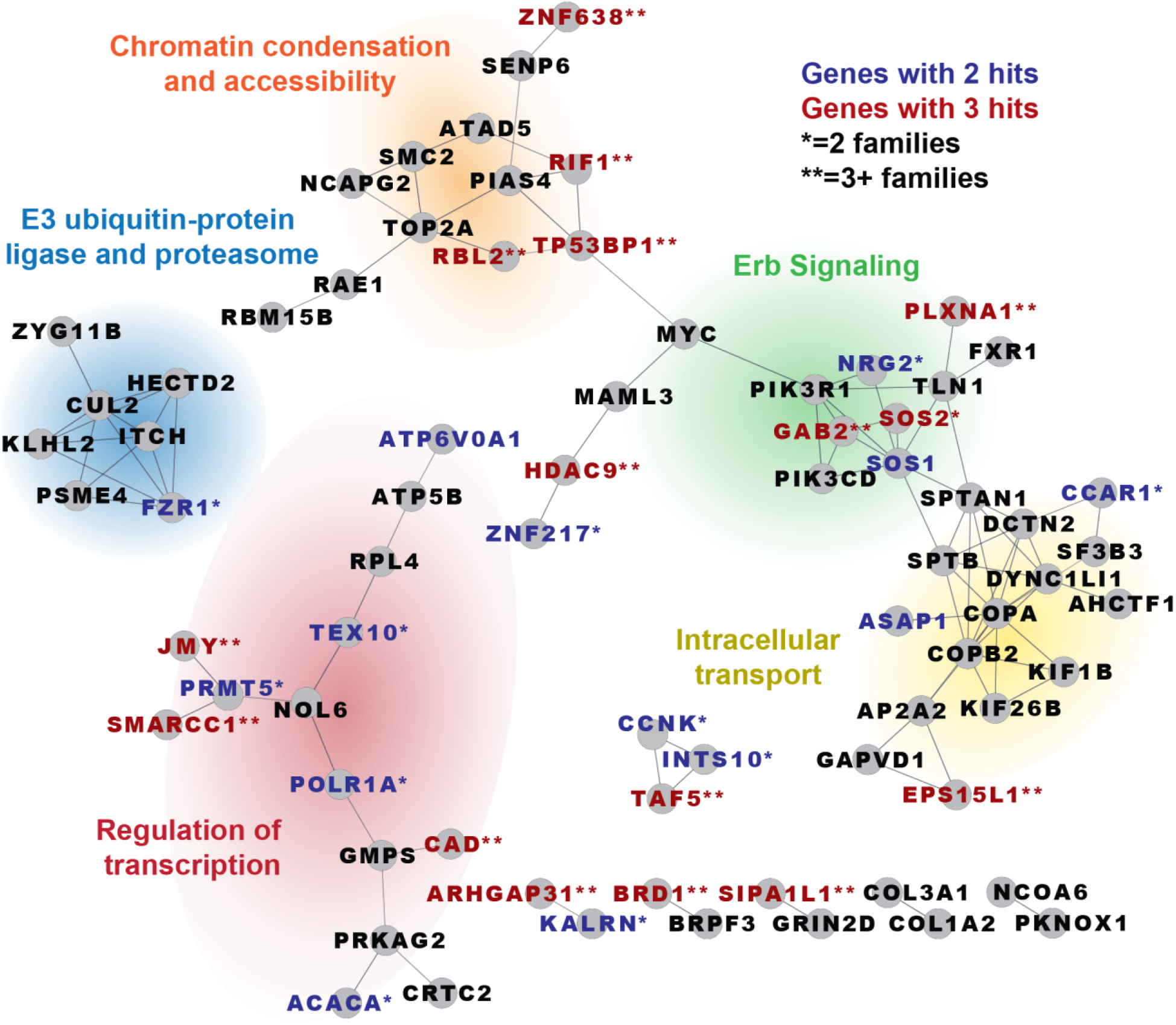
PPI network for autism candidate genes. We identified 163 constrained genes (pLI ≥ 0.99) carrying private LGD mutations that are transmitted only to autism probands based on combined dataset and have not been previously identified as a DNM-enriched ASD gene (Table S9). STRING network shows a significant excess of PPI (p-value = 0.00164). Gene names are colored if observed in 2 (blue) or 3 or more (red) probands and labeled if observed in two independent families (*) or more (**). Families with monozygotic twins (n = 138 in combined) were removed from analysis. Analyses were restricted to regions with at least 20x average coverage in the exomes.

This proband candidate gene set is also enriched for cell-type-specific expression at the early and mid-fetal cortical stages of human brain development (**Fig. S13**). Such an enrichment is not observed if we examine a set of 83 genes in siblings ascertained using the same criteria (not DNM enriched, pLI ≥ 0.99, no private LGD mutations in probands) (**Fig. S13**). If we focus our expression analyses from brain regions to individual cell types in the adult human cortex, we find that our candidate genes are significantly enriched for expression in both excitatory and inhibitory neurons (**Fig. S14**, excitatory p = 4.7e-4, inhibitory p = 5.0e-4) but are not enriched for expression in non-neuronal cell types (**Fig. S14**, p = 0.24) as compared to control sets. However, there is no difference between proband and sibling genes ascertained using the same criteria.

## Private LGD mutations in autism children are evolutionarily younger

Classical population genetics predicts that deleterious variants, such as disease-associated alleles, should be, on average, younger than neutral alleles of the same allele frequency due to purifying selection (*33*). Focusing on children of European ancestry, we applied a genome-wide genealogy method developed by Speidel et al. (*34*) that uses the local ancestry (i.e., linkage disequilibrium) surrounding a single-nucleotide polymorphism (SNP) of interest to construct a coalescent tree and then estimate the generational age of the allele based on the coalescent branch length. To test, we select 101 private LGD variants transmitted only to autism children where none of the 163 candidate genes had been previously associated with ASD. We compare them to a random subset of ~500 private LGD mutations in other genes obtained from both probands and siblings. We estimate the average age of disease-associated LGD mutations to be 2.5 generations and find that these are significantly younger than other classes of private LGD mutations (Fig. 5). For example, other proband-associated LGD mutations outside of the candidate gene set have an estimated median age of 3.7 generations and are significantly older (Mann-Whitney U test, Bonferroni-corrected p-value = 0.0133). Sibling LGD mutations are estimated to be almost two generations older (4.3 generations; Mann-Whitney U test, Bonferroni-corrected p = 0.000255 candidate vs. sibling). As a negative control, we do not observe any difference between the age of private LGD variants in genes outside of the candidate gene set between probands and siblings (Fig. 5, Mann-Whitney U test, p-value = 0.139) or for synonymous or private mutations mapping to intergenic regions (**Figs. S15 and S16**). Since alleles in these candidate genes are carried in unaffected parents, we hypothesize that these variants are under weaker selection than deleterious DNMs but under stronger selection than a neutral allele. Specifically, if we assume mutation-selection balance under an additive (e.g., two-hit; h = 0.5) model (*33*), we can apply gene-specific mutation rates and allele frequencies within the cohort to estimate the median selection coefficient for the 101 private, transmitted LGD variants in probands with European ancestry. We estimate a rather strong selection coefficient of 0.27 (s.d. = 0.24) for private candidate LGD mutations that are transmitted to only autism probands in this study.

**Fig. 5:**
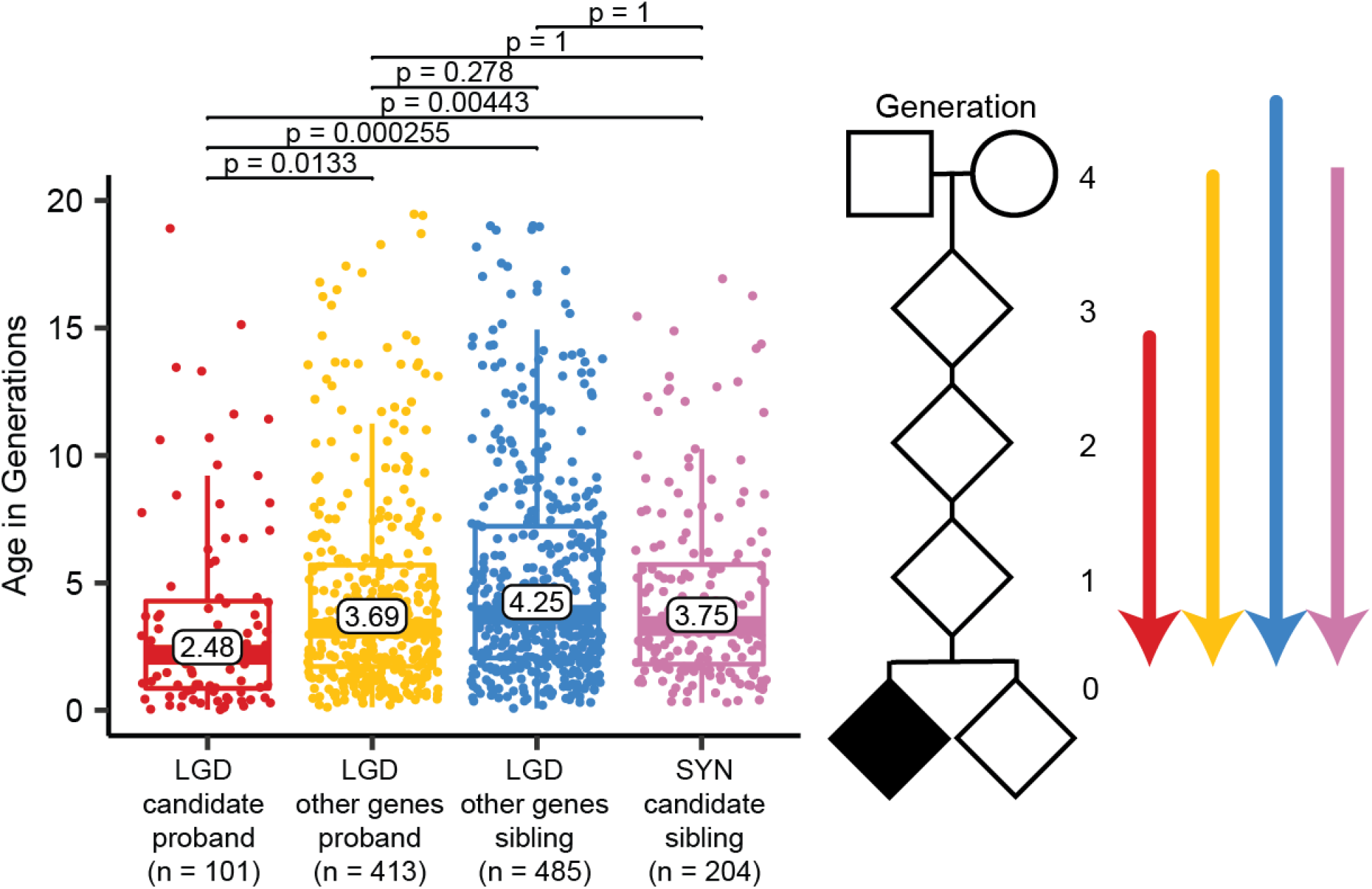
Estimate of allele age. The software Relate was used to estimate the coalescent age (in generations) for private LGD (red) and SYN (blue) mutations in 163 candidate genes and ~500 sites from all remaining genes for European probands (yellow, n = 3,776) and siblings (pink, n = 1,909). P-values are Bonferroni corrected for six tests. Plot was truncated at 20 generations. Data points older than this are included in calculating represented statistics (e.g., boxplots, medians, p-values) but are not visualized. To view all data points, refer to figure S14.

## Discussion

Despite the high heritability of autism, much of the gene discovery in autism research has been driven by studies of *de novo* variation (*2, 7–9, 30*). Our analysis shows that ultra-rare transmitted LGD mutations are not only significantly enriched in autism children but contribute to at least 4.5% of the autism disease risk in the human population. This estimate is in line with other studies (*3, 26*) and suggests that this understudied class of mutation may confer almost as much risk as *de novo* SNVs and indels (6-9% of cases using the same population attributable risk estimator)(*2, 3*). While the burden of private LGD mutations in affected children is higher in multiplex families, both simplex and multiplex families show evidence of transmission disequilibrium for private LGD variants. Interestingly, this effect is significant for simplex families only for genes intolerant to mutation while in multiplex families the effect is larger and significant for genes more tolerant to mutation (Fig. 3B). This may explain why we also observe a significant excess of multiple private LGD mutations in probands as multiple gene disruptions may be required to manifest as autism. We estimate that only about half of the estimated private LGD risk is conferred from genes identified through DNM enrichment studies and that excluding the known DNM-associated risk genes has only a marginal effect on the transmission disequilibrium that we observe.

Most studies focused on identifying risk genes have combined both *de novo* and ultra-rare risk burden. Because a significant fraction of DNM-enriched genes have now already been discovered (*7, 28, 29*), we sought to tease apart these effects by excluding known DNM-enriched genes. To enrich in pathogenicity, we identified a set of 163 candidate genes according to gene constraint (pLI ≥ 0.99) and absence of private LGD mutations in unaffected siblings. Although there has been no reported evidence of DNM enrichment in these genes, we find that several of our candidate genes and gene networks identify pathways previously implicated in autism. For example, we identified three independent private LGD mutations in *HDAC9* that were transmitted exclusively to probands. Pinto et al. identified a transmitted *HDAC9* deletion in a patient with ASD and five additional gene deletions in patients with intellectual disability and schizophrenia (*35*), supporting the role of private, transmitted LGD variants in *HDAC9* in ASD pathogenesis. In addition, several other HDAC genes have also been implicated in ASD, including *HDAC8* (*36*) and *HDAC4* (*37*), and the chromatin remodeling pathway is known to play a key role in autism (*8, 30, 38*). Another gene in our network, *TOP2A*, is part of the topoisomerase gene family thought to be critical in regulating the expression of autism genes (*39*). Although this specific topoisomerase has not yet been routinely observed as disrupted by DNMs in autism families, inhibitors of this gene alter the expression of imprinted genes and the topoisomerase acts by resolving transcription-associated supercoiling of long genes, including ASD-related genes critical for synaptic function.

Additionally, we identified a small network of seven genes involved in the E3 ubiquitin-protein ligase pathway, which has a well-characterized role in autism (*40, 41*). Indeed, there are several genes in this pathway enriched for DNMs in children with autism (*9, 28*), indicating that DNMs and private, transmitted LGD mutations converge on the same pathway but may be hitting distinct sets of genes. An interesting finding from this study has been the discovery of a subnetwork of genes (e.g., dyneins, kinesins, and coatomer subunits) related to vesicular intracellular transport especially between the Golgi and endoplasmic reticulum (Fig. 4). This process is particularly important in the transport of synaptic molecules such as neurexins and neuroligins to the cell surface, endocytic cycling of receptors, and vesicular cargo transport along microtubules (*42–44*). Mutations in related genes in both autosomal recessive and dominant form have been implicated in autism, peripheral neuropathies as well as neurodevelopmental delay where the mutations alter synaptic plasticity and altered morphology of neuronal dendrites and axons. While these associations are exciting, we caution that network and enrichment analyses are often biased toward some of the most well-studied genes and pathways (*45*), and, thus, more than half the genes that failed to associate with a functional networks likely await discovery.

Finally, we report evidence supporting a multi-hit model of autism. Specifically, we find that two or more private truncating mutations in different genes are 50% more likely to occur in autism probands than siblings—a signal consistent with the pathogenicity of this class of variant (Fig. 2A and 3C, **tables S8 and S10**). Interestingly, 56.2% of the 162 children identified in this study (pLI ≥ 0.9) inherit the two mutations from different parents and such transmissions may explain why neither parent is affected. There are other instances of such models reported in ASD ranging from a simple two-hit model (*46*) to an oligogenic model of disease (*4, 10, 11, 16, 46*). Indeed there are several examples of oligogenic mutations playing a role in neurodevelopment, such as the 16p12.1 deletion (*11, 46*), which is often inherited but requires a secondary CNV to reach the genetic liability threshold for disease. Similarly, carrying three or more potentially deleterious DNMs (in the absence of an LGD DNM or large CNV) can be attributed to about 7.3% of autism cases (*4*). In fact, a targeted study of seven genes identified a significant overrepresentation of probands with two or more nonsynonymous variants and suggest that multiple moderate impact events in the same pathway are necessary to cause nonsyndromic forms of autism (*10*). Efforts focusing on patient recontact, not only for the purpose of re-phenotyping families as diagnostic criteria evolve but also for providing additional counseling as novel genetic candidates are identified, will be critical in the task of understanding genotype– phenotype relationships and has already been proposed by others (*47*). Understanding the diversity of genetic etiologies underlying autism as well as their corresponding phenotypic outcomes will be critical for providing accurate disease risk assessments for family planning and genetic counseling.

Taken together, these findings highlight some key considerations for future studies of ASD. Specifically, family composition of the cohort will influence what types of and to what degree different mutation classes contribute to disease risk. This is important for both replication of the findings reported here as well as findings from other groups (*7–9*). The vast majority of autism families characterized by exomes and genomes are simplex in origin and a greater effort must be taken to recruit and characterize multiplex families as part of large-scale sequencing efforts. Additionally, these results highlight the weakness of assuming *de novo* and rare transmitted variants will impact genes in a similar manner (e.g., monogenic and highly penetrant mutations only in constrained genes) (*27, 32, 48*). We find that although *de novo* and private mutations converge on related pathways, there is substantial evidence that suggests these two mutation classes act through different genetic mechanisms and modulate distinct sets of genes in ASD pathogenesis. Our allele age estimates are consistent with the action of strong selection operating on these mutations. Our analysis suggests that the mutations we identify in candidate genes are able to persist for two to three generations before being removed from the gene pool by selection. This is in contrast to most of the LGD mutations associated with *de novo*-enriched genes that are removed from the gene pool almost immediately due to the action of stronger purifying selection.

## ACKNOWLEDGEMENTS

We thank Tonia Brown for assistance in editing this manuscript and Sunday Stray, Mary Eng, James Moore, Hannah Kortbawi, and Anne Thornton from the laboratory of Mary-Claire King for isolation of DNA from whole blood. We thank T. Maniatis and the New York Genome Center for conducting sequencing and initial QC. We are grateful to all of the families at the participating Simons Simplex Collection (SSC) sites, as well as the principal investigators (A. Beaudet, R. Bernier, J. Constantino, E. Cook, E. Fombonne, D. Geschwind, R. Goin-Kochel, E. Hanson, D. Grice, A. Klin, D. Ledbetter, C. Lord, C. Martin, D. Martin, R. Maxim, J. Miles, O. Ousley, K. Pelphrey, B. Peterson, J. Piggot, C. Saulnier, M. State, W. Stone, J. Sutcliffe, C. Walsh, Z. Warren, E. Wijsman). We are grateful to all of the families in SPARK, the SPARK clinical sites, and SPARK staff. We appreciate obtaining access to phenotypic and genetic data on SFARI Base. Approved researchers can obtain the SSC population dataset described in this study (https://www.sfari.org/resource/simons-simplex-collection/) and SPARK population dataset described in this study (https://www.sfari.org/resource/spark/) by applying at https://base.sfari.org. We gratefully acknowledge the resources provided by the Autism Genetic Resource Exchange (AGRE) Consortium and the participating AGRE families. Genomic data for the AGRE cohort provided by iHART, an initiative led by Hartwell Foundation and Directed by D. Wall and D. Geschwind.

## Author contributions

A.B.W. and E.E.E. designed and conceived the study. L.H.W. and M.C.Z. coordinated samples and sequencing for CCDG cohorts. SPARK Consortium coordinated samples and sequencing for SPARK cohort. A.B.W., T.N.T., S.C.M., A.S., T.W., B.P.C., U.S.E., M.B-B., and H.G. called variants and ran QC. K.H. performed Sanger validations. A.B.W., T.N.T., S.C.M., P.H., and A.S. conducted analyses and data interpretation. T.E.B. and A.B.W. performed gene expression analyses. A.B.W., R.A.B. and R.K.E. performed phenotypic analyses. A.B.W. and E.E.E. wrote the manuscript.

## Data and materials availability

The WGS data used in this study is available from the following resources. The AGRE study is available at the Database of Genotypes and Phenotypes (dbGaP) under accession: phs001766. Access to the AGRE WGS data is subject to approval by Autism Speaks and AGRE. All sequencing and phenotype data for the SSC are available through the Simons Foundation for Autism Research Initiative (SFARI) and are available to approved researchers at SFARIbase (http://base.sfari.org, accession IDs: SFARI_SSC_WGS_p, SFARI_SSC_WGS_1, and SFARI_SSC_WGS_2). The genomic and phenotypic data for the SPARK study is available by request from SFARIbase (http://base.sfari.org, globus ID: SFARI:/SPARK/Regeneron/SPARK_Freeze_20190912/). Data from the SAGE study is available at dbGaP under accession: phs001740.v1.p1. Data from the TASC study is available at dbGaP under accession: phs001741. Family-level FreeBayes and GATK VCF files for SAGE, SSC, and TASC are available at dbGaP accession phs001874.v1.p1 and also at SFARIbase under accession: SFARI_SSC_WGS_2a.

## SUPPLEMENTARY MATERIALS

Materials and Methods Figures S1-S17

Tables S1-S13 References 49-62

