## Supplementary text and figures for "Recent ultra-rare inherited mutations identify novel autism candidate risk genes"

**This PDF file includes:**

Materials and Methods
Figs. S1 to S17
Tables S1 to S13 (see supplementary_tables.xlsx)

### Materials and Methods

**Sequencing and quality control of cohorts.** Individuals enrolled in the AGRE, SSC, TASC, and SAGE studies were whole-genome sequenced at the New York Genome Center (NYGC) as part of the Centers for Common Disease Genomics (CCDG, http://ccdg.rutgers.edu/) (**Table 1 and table S1**). This study was approved for sequencing by the local institutional review board (IRB) at the New York Genome Center (Biomedical Research Alliance of New York [BRANY] IRB File # 17-08-26-385). All participants provided informed consent prior to participation in the study (SSC: IRB STUDY00001619 [previously, SAGE: IRB protocol #44219, TASC: STUDY00002514 at the University of Washington). Sequencing was performed on an Illumina HiSeq X Ten platform using 1 ug of DNA and an Illumina PCR-free library protocol. Post-sequencing, the data was processed using the standard pipeline for the CCDG (*49*) and the GRCh38_full_analysis_set_plus_decoy_hla.fa reference genome. Briefly, raw reads were aligned to the GRCh38 reference genome (BWA mem-0.7.15 (*50*)), duplicate reads were marked (Picard v2.5.0), base scores recalibrated (GATK v3.8.0 (*51*)), and indels were realigned (GATK). CRAM quality control (QC) metrics for the SAGE cohort have been previously published (*18*); SSC, TASC, and AGRE QC metrics were determined using Picard WGS metrics, Picard insert-size metrics, and SAMtools (*52*) flagstat. The average sequence depth for SSC, TASC, and AGRE was 34.99 ± 4.09-fold, 33.89 ± 5.46-fold, and 33.03 ± 4.31-fold, respectively. The average insert size for SSC, TASC, and AGRE was 444.9 ± 17.86 bp, 455.1 ± 6.40 bp, and 384.8 ± 26.39 bp, respectively.

Individuals enrolled in the SPARK study were whole-exome sequenced at Regeneron (unpublished) (**Table 1**). Exomes were sequenced to an average coverage of 61.84 ± 14.99-fold. QC analysis included HybridizationMetrics (Picard) and SAMtools flagstat.

**Variant calling.** We called SNVs and indels in families using four different callers: GATK HaplotypeCaller v.3.5.0, FreeBayes v1.1.0, Platypus v0.8.1, and Strelka2 v2.9.2. In addition, multi-nucleotide variants were called using FreeBayes and Platypus. Post-calling, BCFtools (version 1.3.1) norm was used to left-align and normalize indels. We partitioned the genome into the high-quality regions, consisting of unique space as well as ancient repeats and the recent repeat regions, which consisted of repeats <10% diverged from the consensus in RepeatMasker. Variants were only assessed in high-quality portions of the genome and recent repeat region variants were removed from the study.

**Kinship and sample redundancy.** All samples from both the discovery and validation cohorts were merged together and kinship coefficients were calculated with KING (v1.4) (*53*). Samples with kinship coefficients that did not match their reported relationship were identified as potential sample swaps or contamination and were either removed or, when possible, their relationships were corrected. Samples with kinship coefficients greater than 0.35 were identified as potential sample duplicates (**Table S13**). We first checked whether potential duplicates were known monozygotic twin pairs or known duplicates within a cohort (some individuals had both blood and cell line DNA sequenced for QC purposes). We retained one sample from each of the known duplicate pairs, preferentially retaining the sample generated from blood DNA when possible and randomly selecting the retained sample if there was no difference in DNA source.

The remaining duplicates, which represented samples that were sequenced as part of multiple cohorts were retained for one and only one of the cohorts according to the following prioritization scheme: 1) sample was sequenced as part of an SSC family; 2) the sample was from a complete family, their DNA was from blood, and was WGS; 3) the sample was from a complete family and sample was WGS; or 4) the sample was from a complete family and contained unaffected siblings. Families with twins were retained for private variant discovery but were excluded from all statistical analyses.

**Principal Component Analysis (PCA).** In addition to our discovery and validation cohorts, we included two reference cohorts, 1000 Genomes Project (1KG, 20140818 release) (*24*) and Simons Genome Diversity Project (SGDP, available at NCBI under BioProject ID PRJNA522307) (*54*), in our PCA. Each cohort was cleaned separately (described below), merged together, and then cleaned additionally. Reference data from the SGDP and 1KG was prepared for PCA by left normalizing variants with BCFtools v1.9, followed by filtering for individual missingness (<10% missing genotypes within an individual), SNP missingness (<50% missing genotypes across a SNP), minor allele frequency (>5%), and linkage disequilibrium pruning with PLINK 1.90 (*55*). Sites were then converted from hg19 to GRCh38 using UCSC liftOver. Since the 1KG data was generated with a lower density SNP array than SPARK, the remaining 1KG sites were the only sites considered in the remaining cohorts. The SSC, SAGE, TASC, and AGRE samples were prepared using GATK joint-genotype files provided by the NYGC and then iteratively merged together within the respective cohort (most cohorts had to be processed in multiple batches). Each joint-genotype file was prepared as described above. The SPARK samples were prepared using the joint-genotype generated by Regeneron using Illumina

InfiniumCoreExome-24_v1.1 array data. Intensity data files were processed using Illumina Genome Studio Software. Since this data was already in plink format, it did not undergo additional processing. Prior to merging all six cohorts together, the 1KG target sites were extracted from each cohort. After merging, the combined autism and reference cohorts were filtered for genotype missingness within the individual and SNP (both < 5%). Finally, the data is input into EIGENSTRAT v5.0.1 for PCA. The results of this analysis are summarized in **Fig. S17**.

**ADMIXTURE and ancestry assignment.** The files we used for PCA input were split by reference cohort (SGDP and 1KG) and discovery cohort (SSC, SAGE, TASC, and AGRE) and filtered for sites present in the reference cohort and individual-level missingness a second time. We ran the software ADMIXTURE v1.3.0 (*56*) with 10-fold cross validation (CV) on our reference cohort of 1,964 unrelated individuals to determine the optimal value for the K parameter. We found that K=10 resulted in the smallest CV error (**Figure S2C**); however, there is little difference in CV error for values of K between 8 and 14 and we recognize that a lower value of K would result in similar population assignments. We assessed the quality of our inferences for our reference cohort by visualizing the proportion of ancestry from each cluster for a random subset of 15 individuals from each known population (250 individuals total).

Due to the underlying relationships between individuals in our autism cohort, we chose to use the allele frequencies learned by ADMIXTURE from our reference cohort to assess the ancestry of our discovery cohort in a supervised manner by using projection with ADMIXTURE. We assigned each individual to the cluster that contributed the largest proportion of ancestry and then grouped clusters into six super populations (EUR = European, AFR = African, EAS = East Asian, SAS = South Asian, AMR = Amerindian, OCN = Oceanian) according to membership of known populations from the reference cohort (**Figure S2A**). We were unable to assign ancestry to 1.01% of our discovery cohort due to missing data in the joint-genotype files and find that the majority of our cohort (85.9%) is European (**Fig. S2B and table S4**).

***De novo* mutation (DNM) calls.** DNMs in the SSC, SAGE, and TASC cohorts were called using a custom pipeline. DNMs were not called in AGRE because DNA for most samples in this cohort was derived from cell lines, which are prone to introducing artifacts in DNM analyses. First, variants that were *de novo* based on genotype (father and mother genotypes were equal to 0/0 and the genotype in the child was 0/1 or 1/1) were retained for further assessment. Second, variants from Platypus with a filter of LowGQX or NoPassedVariantGTs were removed and Strelka2 variants had to have the filter field equal to PASS. Third, variants needed to have the support of at least two of the four callers. Fourth, variants were regenotyped with FreeBayes using default settings and needed to remain as *de novo*. Fifth, variants in a homopolymer A or T of length 10 or greater were removed. Sixth, we removed all variants in low-complexity regions, recent repeats, or centromeres. Finally, we applied the following sample level filters: the father alternate allele count = 0, mother alternate allele count = 0, child allele balance >0.25, father depth >9, mother depth >9, child depth >9, and either child genotype quality (GQ) >20 (GATK) or sum of quality of the alternate observations (QA) >20 (FreeBayes). For variants on the X chromosome we separately considered variants in the pseudoautosomal regions (chrX:10000-2781479, chrX:155701382-156030895) and the X/Y duplicatively transposed region (chrX:89201803-93120510).

We performed random Sanger validation and combined this data with published validations to look at a total of 3,233 sites in a conditional inference analysis (**Table S2**). The metrics we included in this analysis included 1) the mer150 mappability, which we calculated on build 38 of the human genome using a workflow originally designed as part of the ENCODE project; 2) the average mapping quality of the read +/- 100 base pairs (bp) around the variant in the child; 3) the average mismatch in the reads +/- 25 bp around the variant in the child; and 4) the callers that supported the event as *de novo*. Based on this analysis, the final dataset for *de novo* SNVs and indels were sites that either had the support of all four callers or were supported by three callers and had an average mapping quality greater than 57 for the reads in the 100 bp region around the variant. For the multi-nucleotide variants, we also inspected all sites using SAMtools tview and the sites had to have visual inspection support of *de novo* status and an average mapping quality greater than 57 for the reads +/- 100 bp around the variant. We estimate our validation rate in this dataset at 99.5% and our false negative rate at 3.5%. In addition, we removed samples that were statistically defined as outliers, in terms of *de novo* counts, based on the boxplot function in R.

**Private SNV calls.** Each cohort was assessed separately to identify ultra-rare, inherited variants using a custom pipeline. Briefly, SNVs and indels were called using FreeBayes (v1.1.0) and GATK on a per-family basis. Sites were left-aligned, normalized, and multiallelic sites were split into separate lines using BCFtools v1.9. Sites from the two callers were merged using GATK CombineVariants. To ensure a high level of specificity, we counted all alleles in the parent population that were present in the union set of the two callers and passed the following QC filters: 1) site quality score (QUAL) > 50, and 2) read depth (DP) ≥ 10 for genomes and DP ≥ 20 for exomes. We used slightly different DP filters for the exome and genome data to account for differences in sequencing depth between the two sequencing platforms. All sites that were heterozygous and observed only once in the parent population were designated as candidate private variants (cohort-level parental frequency ≤ $7e^{-5}$; approximate equivalent ExAC frequency ≤ $2.5e^{-5}$).

The set of private variants for each cohort was comprised of candidate private variants that were present in the intersection set of GATK and FreeBayes and did not violate the rules of Mendelian inheritance. We annotated variants using SnpEff v4.3t with gene and transcript information (GRCh38.86), predicted effect of the variant on the transcript, ExAC (r0.3, non-neuropsych subset) lifted over to GRCh38 using the UCSC liftOver tool, and dbSNP (v150). Finally, variants were filtered against recent repeats (see DNM methods for details), low-complexity regions, centromeres and gaps, and pseudoautosomal regions (hg38 chrY:10,000-2,781,479, chrY:56,887,902-57,217,415, chrX:10,000-2,781,479, chrX:155,701,382-156,030,895) using BEDTools v2.24.0.

The set of private variants from each cohort was compared to all variants observed in the other cohorts. Candidate private variants that were not observed in any other cohorts were retained for our final set of private variants. For example, the discovery cohort private variants are comprised of sites unique to one parent across only the WGS cohorts, whereas the combined set private variants are comprised of sites unique to one parent across both the WES and WGS cohorts. When combining the WES and WGS cohorts, we only included regions with an average coverage of 20-fold in the exomes.

**Transmission bias and burden.** We partition mutations from protein-encoding regions of the genome into three classes: 1) likely gene disrupting, which we define as any mutation that introduces a stop codon, ablates a stop, changes the frame of the open reading frame, or introduces a change at a predicted splice donor or splice acceptor site, 2) missense, which is any mutation that causes an amino acid change, or 3) synonymous, or any mutation that results in no amino acid change. We quantified the number of private transmitted variants observed in probands and unaffected siblings by gene set and variant type and compared the proportion of carriers using both a Fisher's exact test and logistic regression (one model for each variant type and pLI threshold). For the DNM-enriched gene set analyses, we compared the proportion carriers between probands and siblings using Fisher's exact test and logistic regression. Multi-hit analyses and simplex versus multiplex analyses were conducted using a Fisher's exact test to compare the proportion of individuals carrying two or more hits in probands and siblings. We applied Bonferroni and false discovery rate corrections to all p-values using the R function p.adjust for each analysis.

**Population attributable risk (PAR).** PARs were calculated using the following formula published by Cole and MacMahon (*57*). Our calculations assume that siblings are representative of the general population. Even if siblings are sub-threshold for ASD, these estimates would serve as a lower bound for PAR.

$P_e$ *= proportion of population (controls) exposed*

*RR = relative risk or ratio of the risk between the exposed and unexposed*

$$PAR\ (\%) = \frac{P_e \times (RR-1)}{(1+ P_e \times (RR-1))}$$

**Allele age estimation.** We estimate the age of a private, transmitted variant at a site of interest using the software Relate (*34*). In short, Relate reconstructs the local genealogy of the region of interest using a scalable computation, which guarantees the inferred genealogy exactly producing the observed data under the infinite-sites model and, thus, can effectively apply to datasets with thousands of individuals while taking into account recombination. Mutations are then mapped onto the branches of the resulting local tree to estimate mutation age. Each private, transmitted variant is annotated as LGD, synonymous, or intergenic as described above and then classified into one of the three datasets for candidate gene, affected proband, and unaffected sibling depending on the carrier of the private variant. To reduce the computational burden for the inferences for synonymous and intergenic sets, we divided the genome into 100,000 bp windows and randomly selected a locus of interest from up to 550 windows (up to 25 windows per chromosome). We removed sites where the derived allele could not be determined.

For a site of interest, we first generated phased haplotypes for the 100,000 bp region surrounding the site of interest for all samples using BEAGLE (v5.1) (*58*) without imputation. For our analysis, we included only one individual from each family and removed samples that are likely related (see Methods for kinship above). To model recombination wherever applicable, we used the HapMap genetic map. Ancestral and derived states for individual sites are based on the sequences downloaded from Ensembl. To ensure the quality of the genotypes, we masked sites that are within segmental duplications, low-complexity, and repeat masker sequences (see Methods for private SNV calls above). Relate outputs two age estimates: The lower and upper ages represent the ages of the coalescence events below and above the mutation of interest, respectively. We determine the age of a given variant by taking the average of the two estimates. Note that we recognize the presence of natural selection at sites where deleterious mutations occur would affect the inference of allele age, and thus, the age estimates for deleterious alleles inferred in this study are overestimated and can be deemed as an upper bound for their ages. We compare the distributions of allele age among different datasets using the Mann-Whitney U test and p-values are all Bonferroni-corrected using p.adjust. Potential caveats, such as phasing errors and cryptic relatedness, might affect the individual age estimates but are expected to have limited impacts on the observation of differences among different variant sets because the same procedure was applied to individual sets.

**Selection coefficient estimation.** We apply the classic mutation-selection balance model to estimate selection coefficients for the 101 private, transmitted variants in the candidate gene set. Our rationale is that because these variants are predominantly recent (on the order of 1/s) (*33*), selection must act relatively strong on these variants, and thus the presence of these variants are primarily due to mutation. Assuming the multi-hit, additive model (h=0.5), the individual selection coefficient (s) can be approximated as mu/qh, where mu is gene-specific mutation rate, h is the dominance coefficient, and q is the observed allele frequency in the entire population sample.

**Gene expression analyses.** Cell-type specific expression analyses (CSEA) were conducted using the CSEA tool (*59*). Candidate gene from probands and siblings were uploaded to the available web server for CSEA across brain regions and development in humans. Gene expression was identified in a published set of transcriptomically defined cell types in the human temporal cortex

(*60*). Gene sets were tested for enriched expression in three broad cell classes – inhibitory neurons, excitatory neurons, and non-neuronal cells – by counting the number of cell types within each class that expressed each gene with average counts per million (CPM) greater than 1. For each gene set, the number of cell types with expression were calculated, and gene sets were visually compared by plotting cumulative distributions. For each cell class, Wilcoxon rank-sum tests were used to identify statistically significant differences in the number of cell types with expression for ASD and control gene sets. P-values were adjusted for multiple comparisons using Bonferroni correction.

**PPI network analysis.** We used the STRING database (STRING) v11 to perform PPI network analyses via Cytoscape v3.7.2 (*61, 62*). We used the multiple protein input option with all default settings except we required interactions to be limited to those which were high confidence (0.700). Disconnected nodes were hidden from the network output (but not from enrichment analyses). In addition, we also used STRING to calculate network statistics and run functional enrichment analyses from the Gene Ontology resource, KEGG, and the Reactome pathway database to identify shared functions across the full set of genes and three subnetworks. A subnetwork was identified as any group of genes that contained at least five genes. The genes *KIAA0430* and *ATP5B* are also known as *MARF1* and *ATP5F1B*, respectively.

All statistics were calculated using R versions 3.5.1 and 3.5.2.

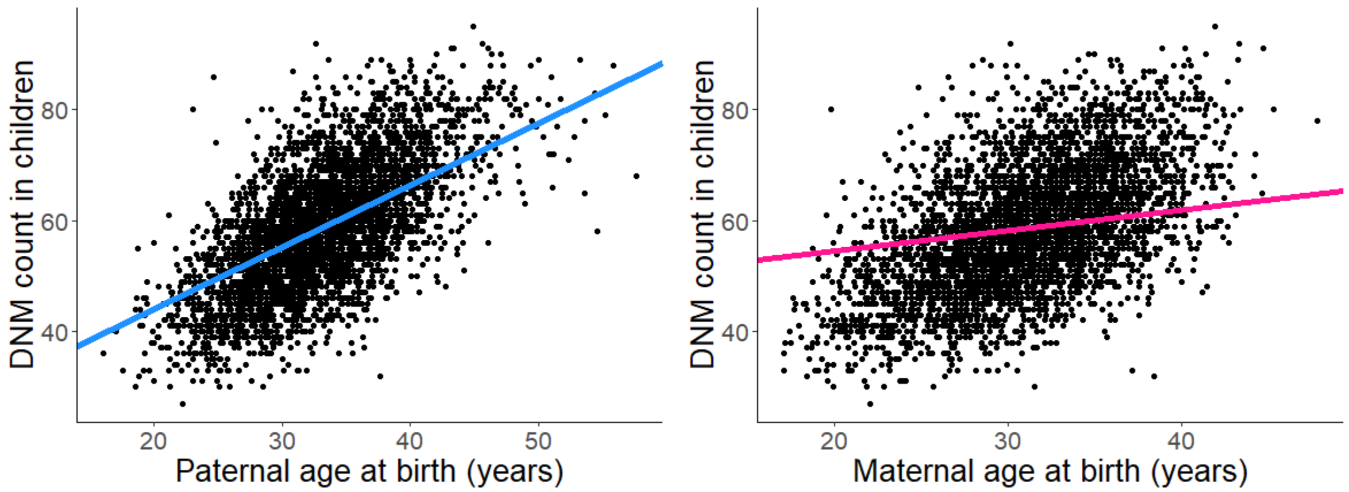


**Fig. S1: Relationship between parental age and DNM counts in children.** We observe a significant correlation between parental age and DNM counts in children. As expected, fathers contribute, on average, more DNMs per year of age than mothers (1.11 vs. 0.37).

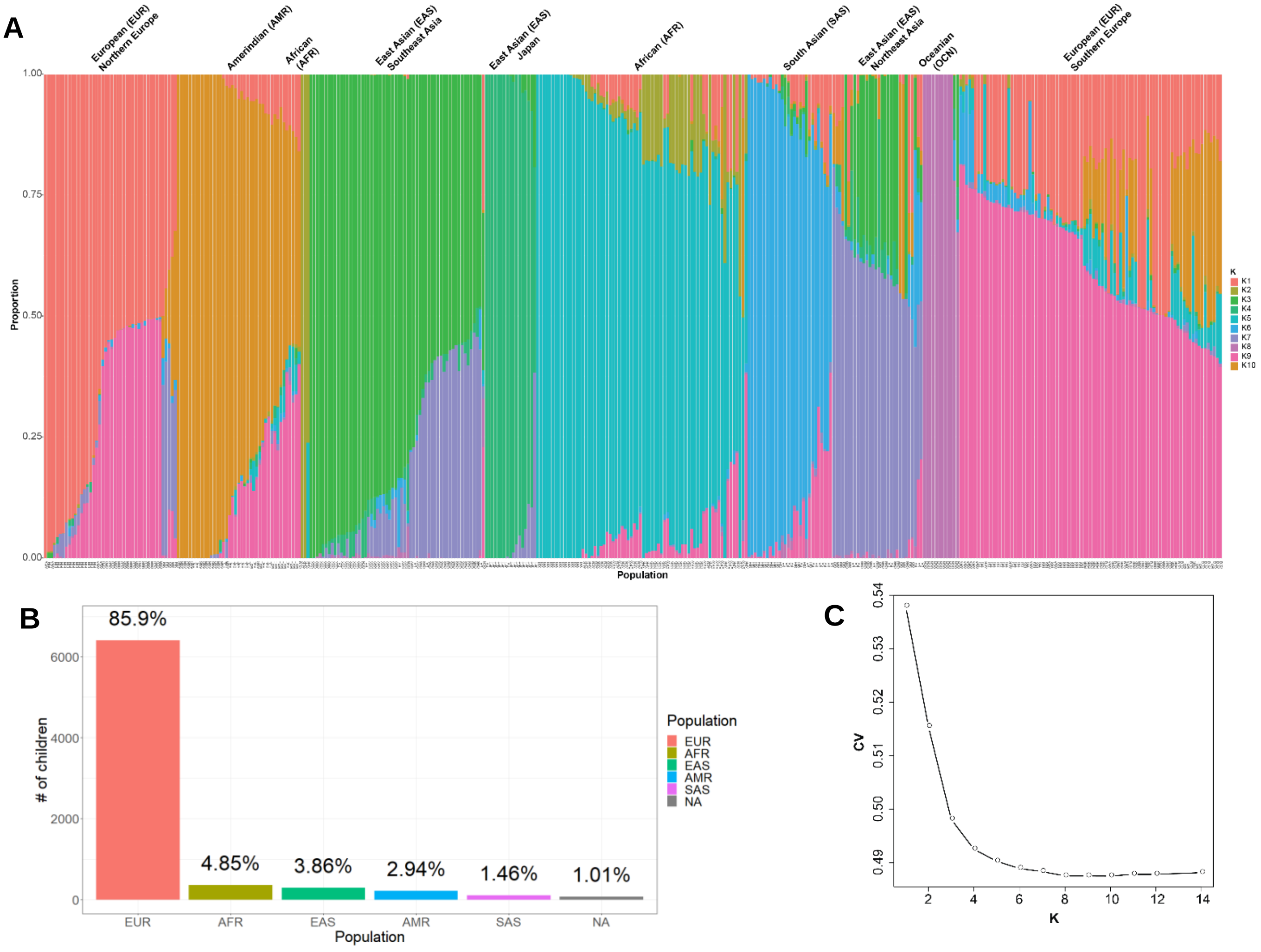


**Fig. S2: Ancestry inference for diversity panel and autism family samples.** A) Plot of ancestry proportions as estimated by ADMIXTURE for 15 randomly sampled individuals from the SGDP and 1000 Genomes Project populations from each reported population. Known population for each individual is labeled on the x-axis and likely geographic origin is noted above each cluster. B) Ancestry assignments for our discovery population. Ancestry for each individual was assigned according to the population with the largest proportion in that individual. C) Cross-validation (CV) error across different K for ADMIXTURE.

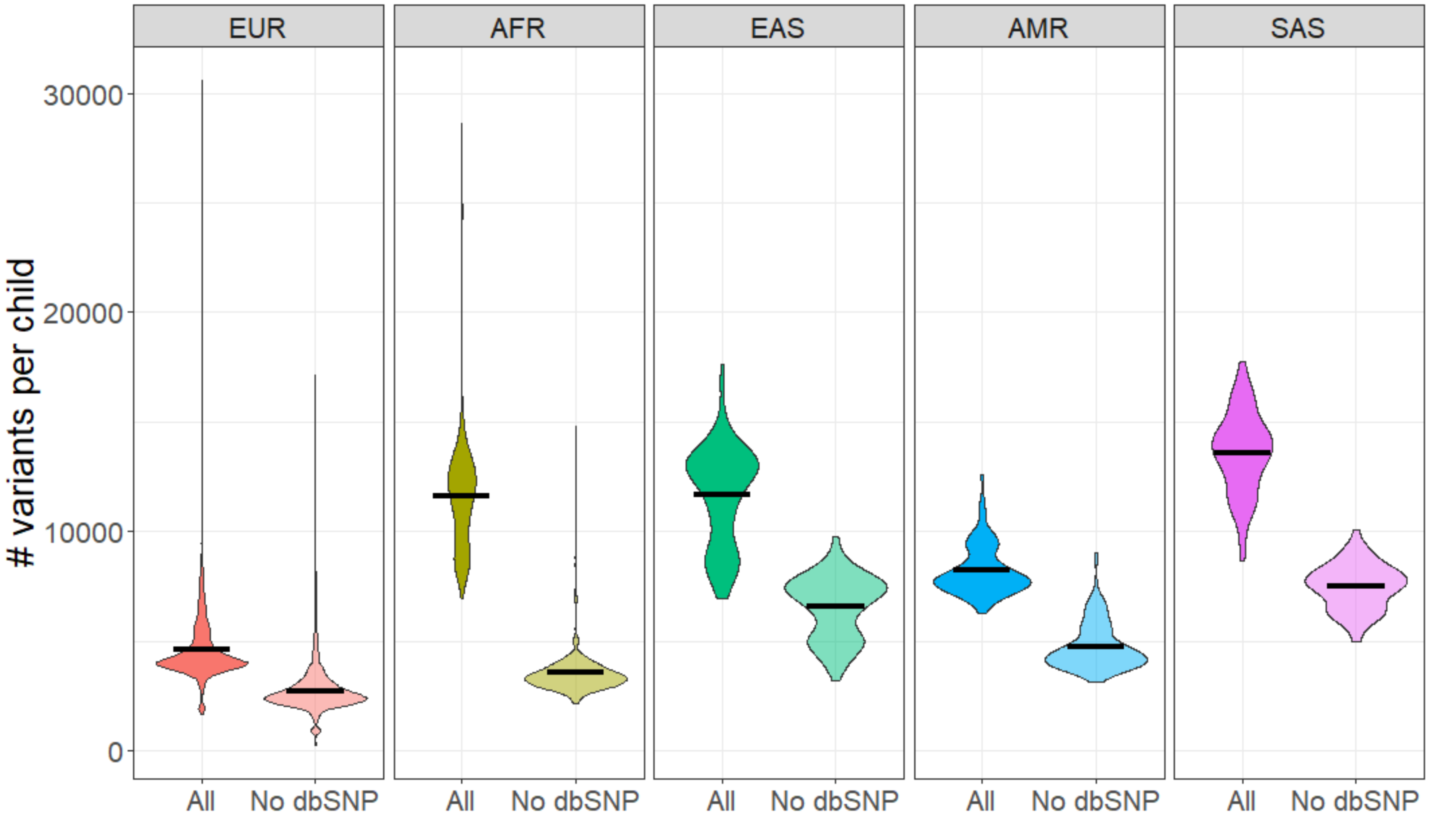


**Fig. S3: Private, transmitted variant counts.** Variant counts per child grouped by ancestry (EUR=European (n = 5,685), AFR=African (n = 290), EAS=East Asian descent (n = 252), AMR=Amerindian (n = 193), SAS=South Asian (n = 103)) before (All) and after (No dbSNP) filtering with dbSNPv150. Excess of private variants is partially but not fully resolved after excluding sites observed in dbSNP. We were unable to assign ancestry to one of these five population groups for 74 of the children in this study.

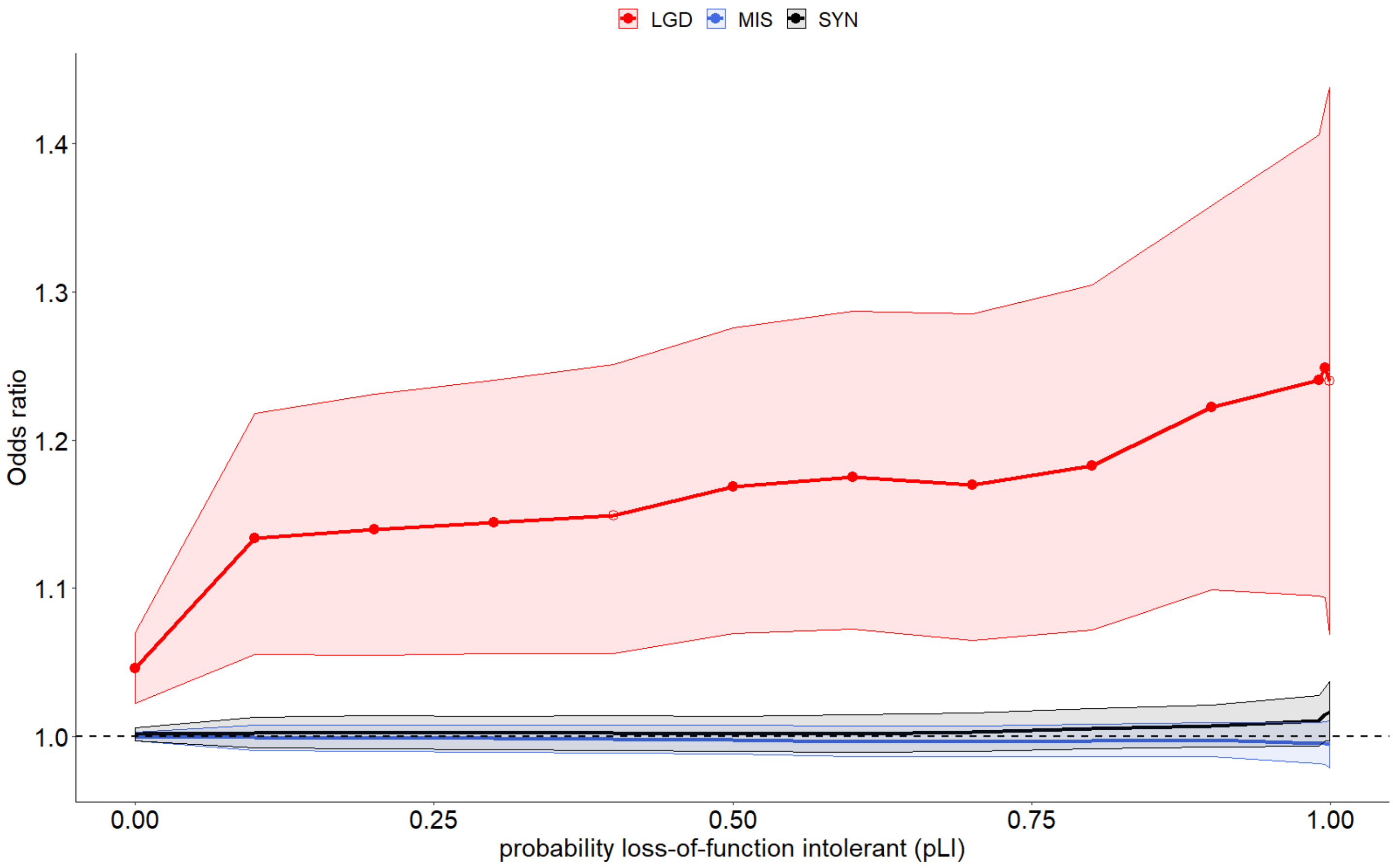


**Fig. S4: Patterns of private mutation burden with gene constraint in EUR subset.** Burden of private LGD mutations in probands increases with gene constraint in EUR subset of discovery cohort (n = 3,636 probands and 1,873 siblings). Excludes families with monozygotic twins. Filled circles indicate Bonferroni-corrected p-values < 0.05 (42 tests). Unfilled circles indicated uncorrected p-value < 0.05.

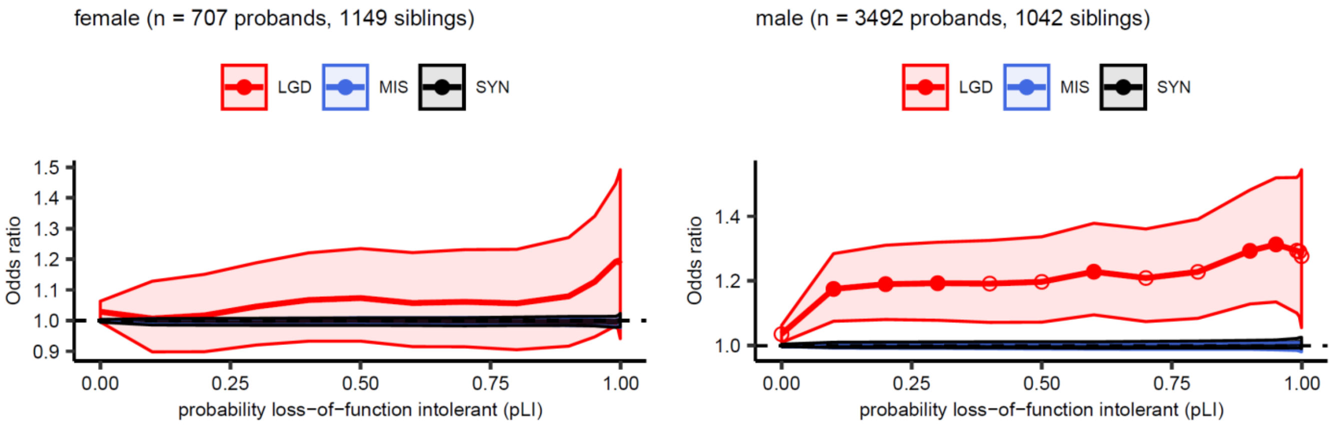


**Fig. S5: Patterns of private mutation burden with gene constraint by sex.** Burden of private LGD mutations in probands increases with gene constraint in males and females. Filled circles indicate Bonferroni-corrected p-values < 0.05 (84 tests). Unfilled circles indicated uncorrected p-value < 0.05.

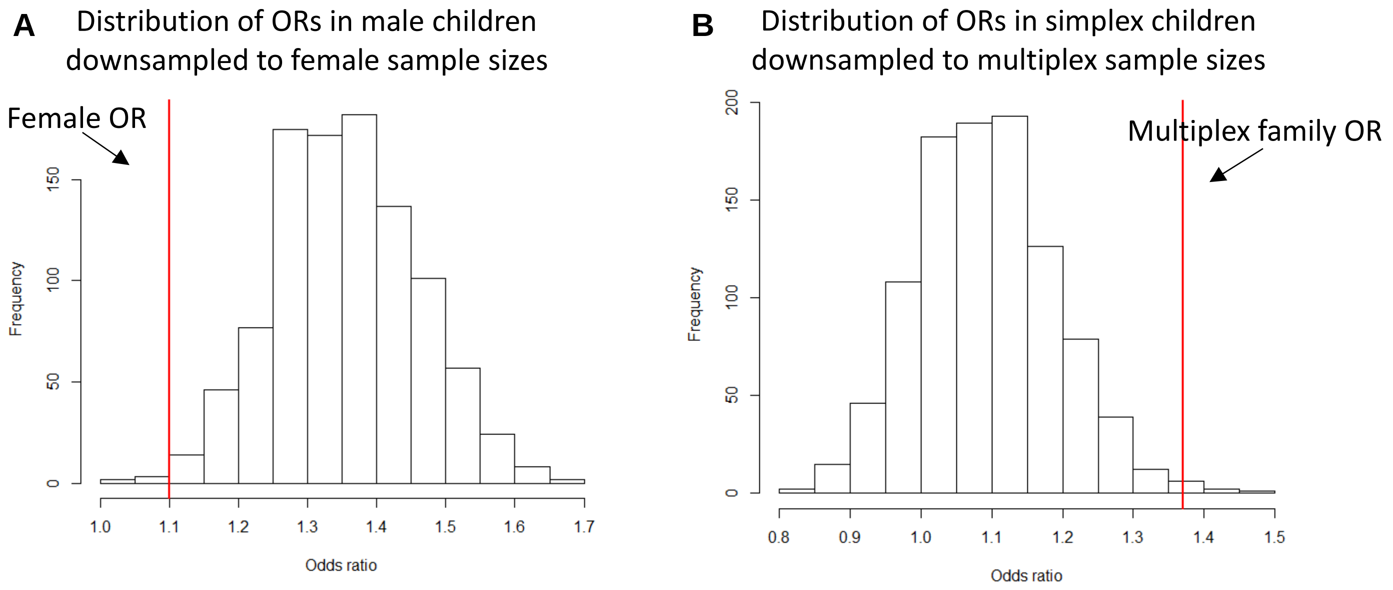


**Fig. S6: Permutation tests for estimated burden in probands.** A) Male permutation tests were performed by randomly sampling 707 male probands to match the sample size of the female probands. Since there are ~100 fewer male siblings as compared to female siblings, all male siblings were used (i.e., no downsampling for male siblings occurred) and then the burden of private LGD mutations in probands vs. siblings in genes with pLI ≥ 0.9 was calculated using a Fisher's exact test. B) Simplex tests were performed by randomly sampling 2,691 probands and 533 siblings to match the sample sizes observed for the multiplex families and then calculated the burden (OR) of private LGDs in probands vs. siblings in genes with pLI ≥ 0.1 using a Fisher's exact test. Red lines indicate the observed OR for females and multiplex, respectively. We performed 1,000 permutations of each analysis.

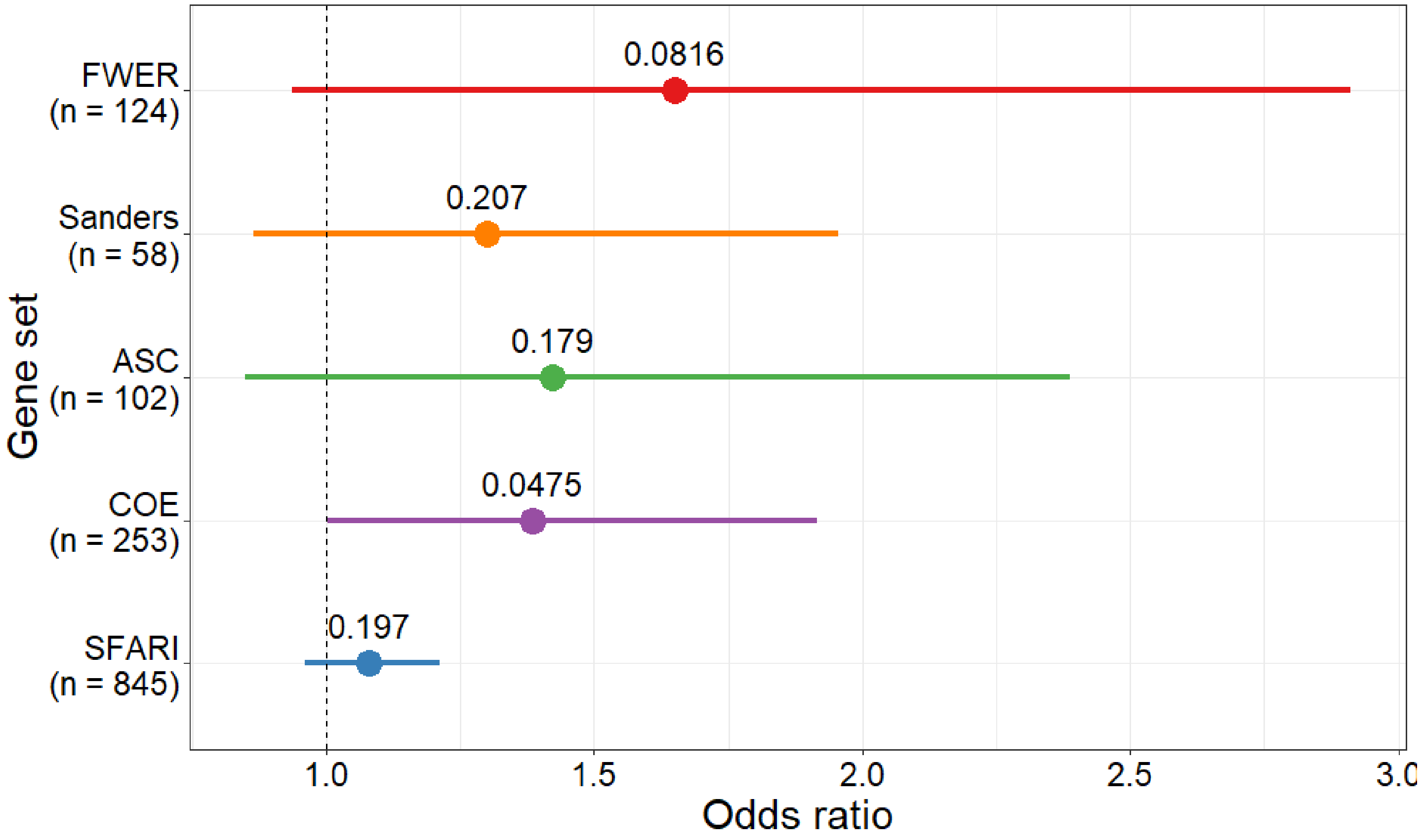


**Fig. S7: Private LGD mutation burden across five DNM-enriched gene sets in EUR subset.** EUR subset of discovery cohort is comprised of 3,636 probands and 1,873 siblings. Analysis excludes families with monozygotic twins. Reported p-values are nominal.

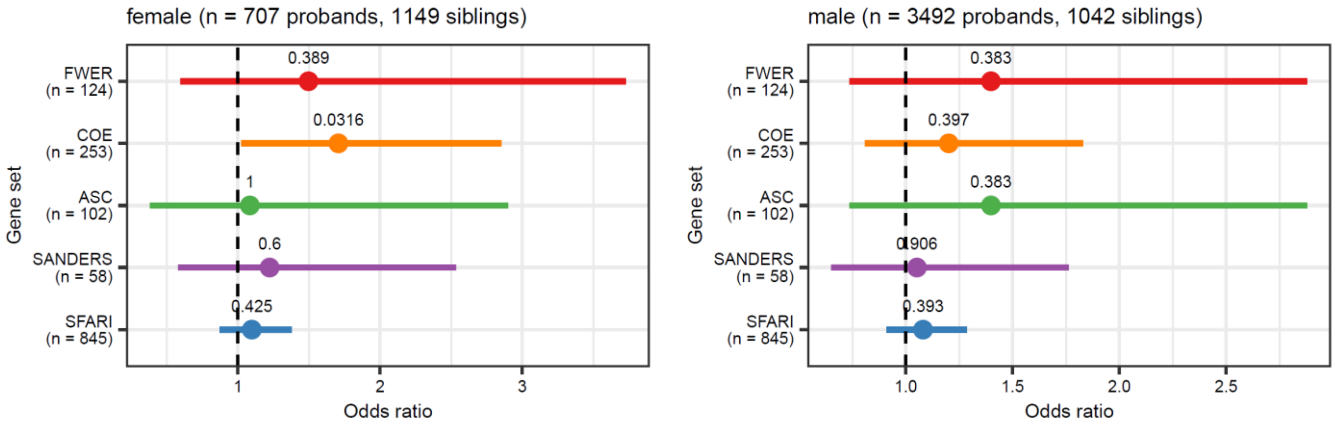


**Fig. S8: Private LGD mutation burden across five DNM-enriched gene sets in males and females.** No gene sets reach Bonferroni-corrected significance for ten tests.

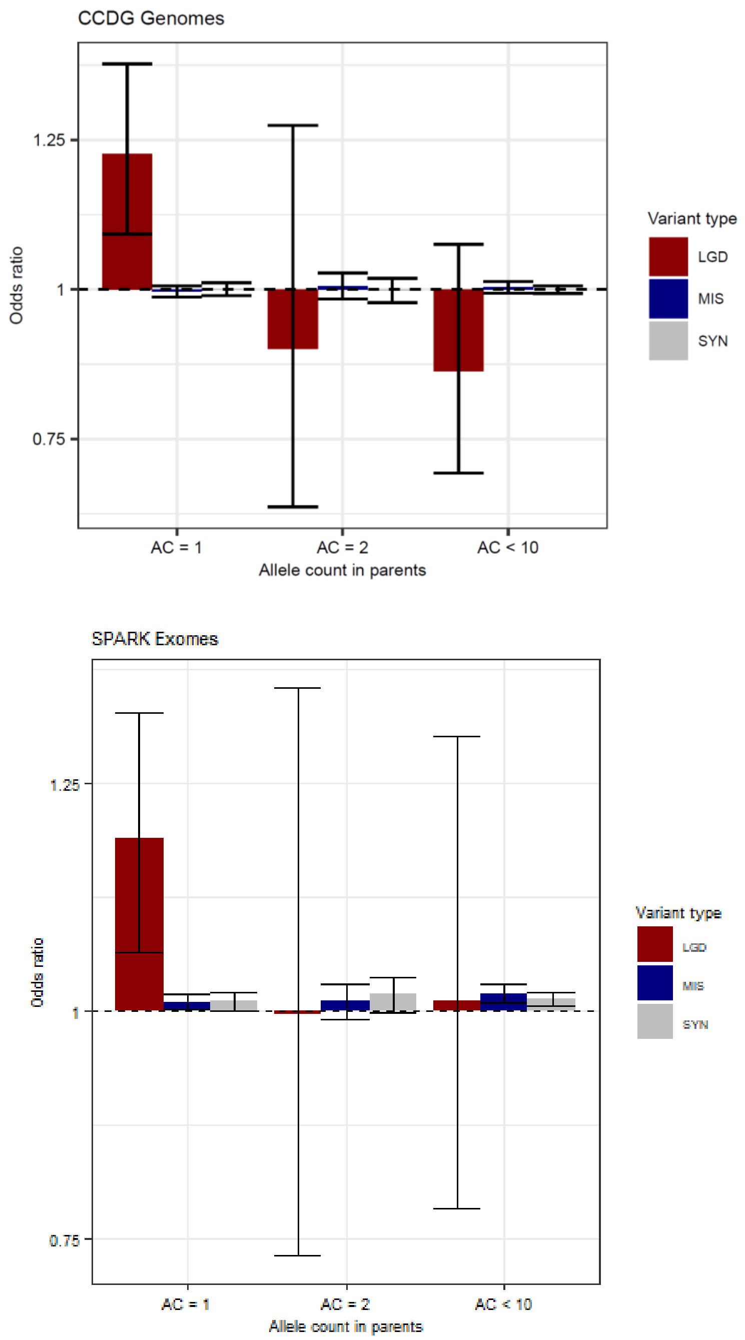


**Fig. S9: LGD burden in probands isolated to private variants.** Burden of LGD, MIS and SYN variants were compared in probands vs. siblings at three allele count bins using a Fisher's exact test for pLI $\geq 0.1$. Analyses exclude families with monozygotic twins.

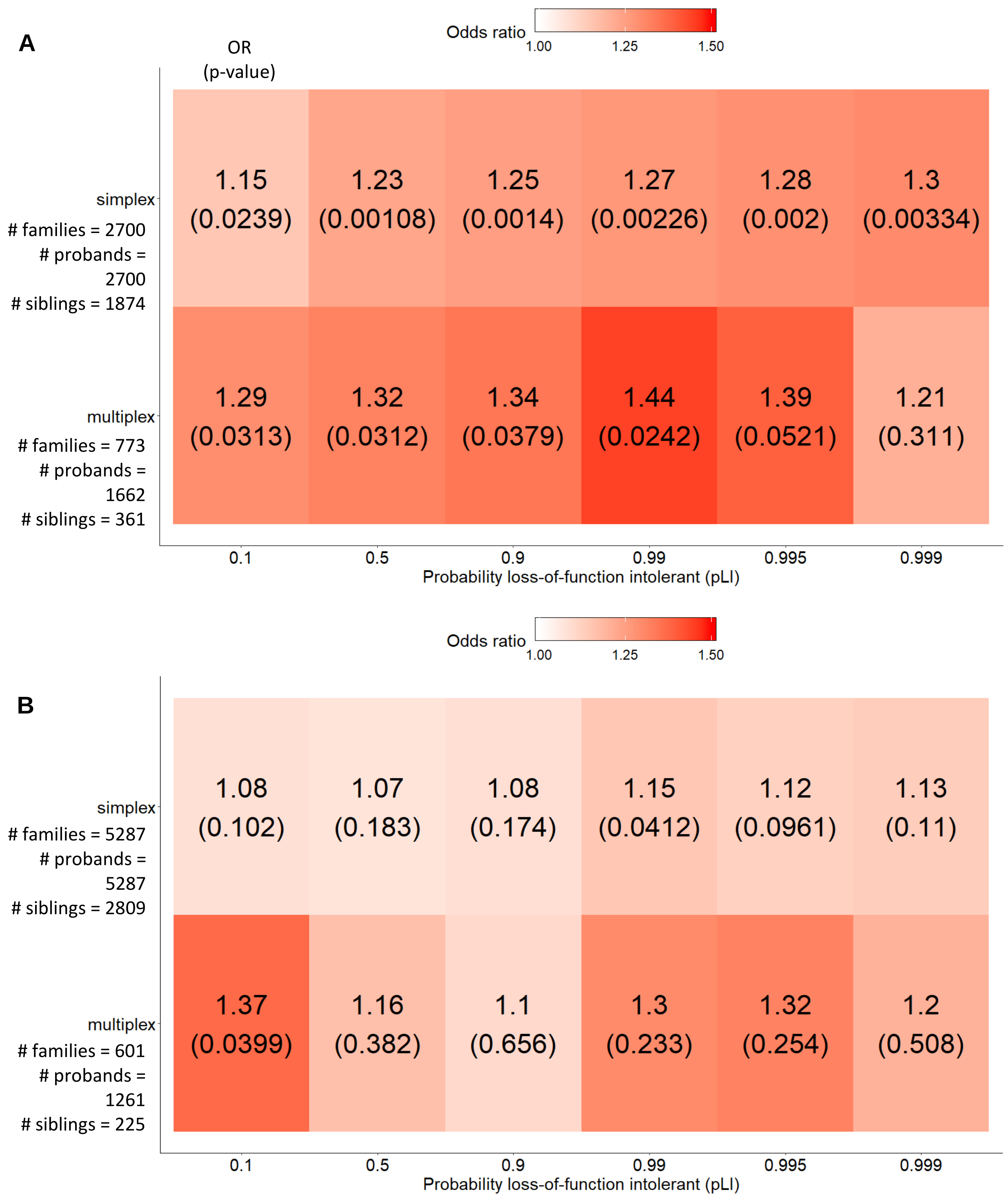


**Fig. S10: Burden of private LGDs in simplex and multiplex families.** A) Our discovery cohort and B) our replication cohort. Excludes families with monozygotic twins. Reported p-values are Bonferroni corrected for nine tests.

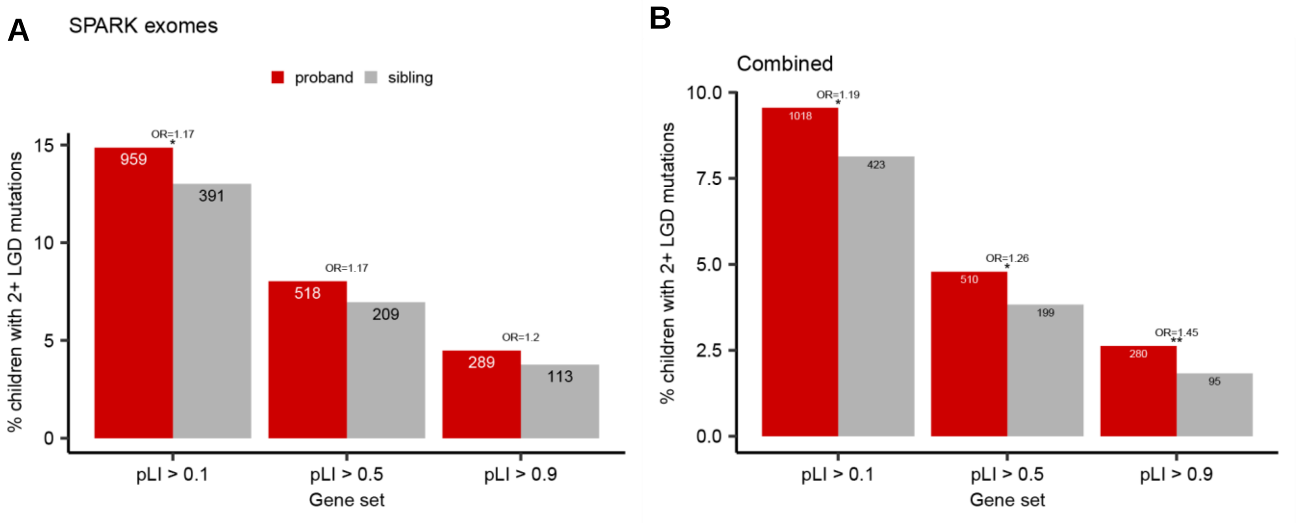


**Fig. S11: Burden of 2+ private LGD mutations.** A) Replication cohort (n = 6,539 probands, 3,034 siblings) and B) combined discovery and replication cohorts (n = 10,638 probands, 5188). Probands are enriched for multiple private, transmitted LGD mutations as compared to siblings across increasing thresholds of pLI. Families with monozygotic twins (n = 75 in discovery, n = 63 in replication, and n = 138 in combined) were removed from analysis. For the combined set, variants were restricted to regions with at least 20x average coverage in the exomes. Significance stars reflect Bonferroni-corrected p-values for three tests.

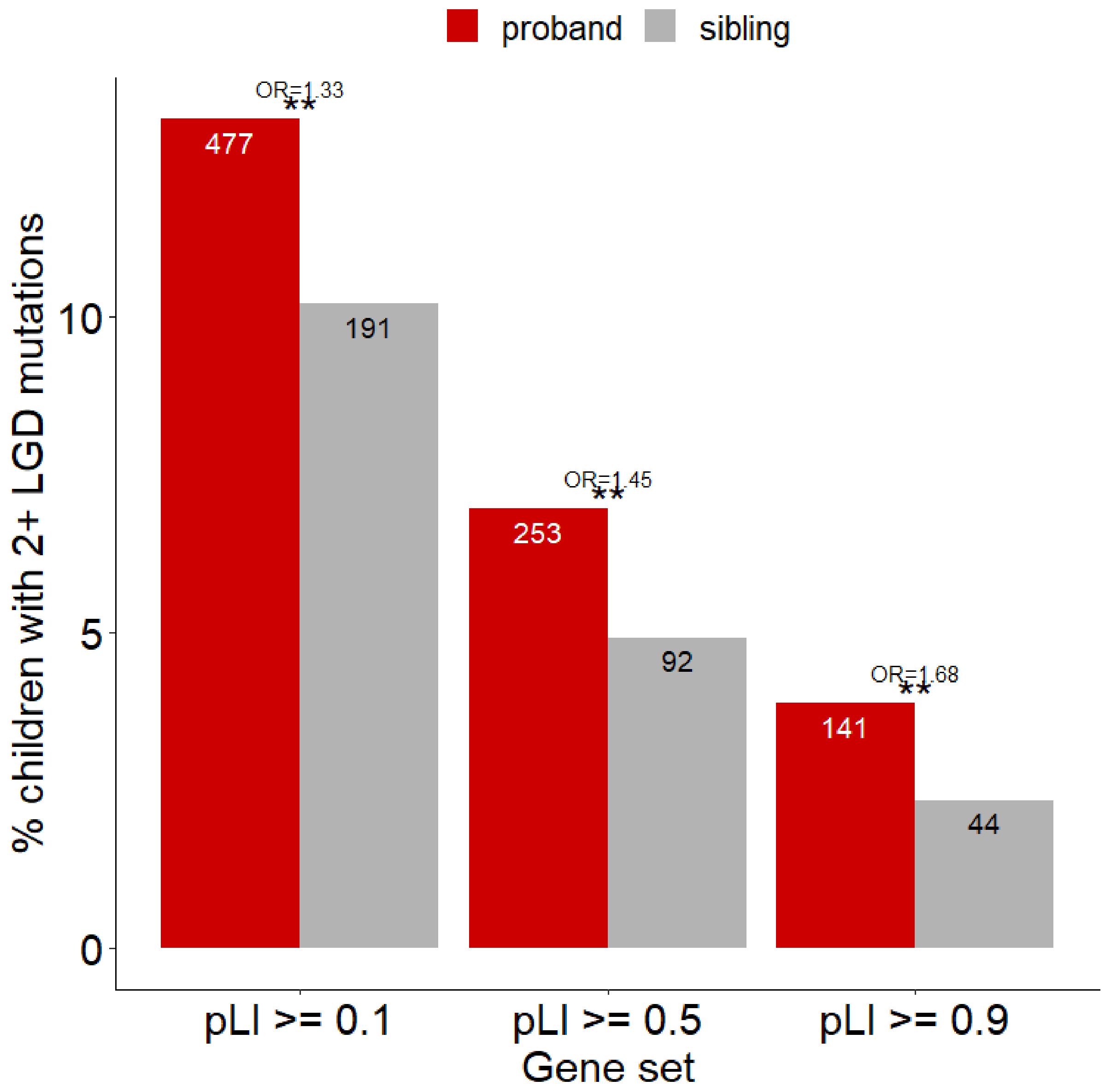


**Fig. S12: Burden of 2+ LGD mutations in the EUR subset.** EUR subset of the discovery cohort is comprised of 3,636 probands and 1,873 siblings. Excludes families with monozygotic twins.

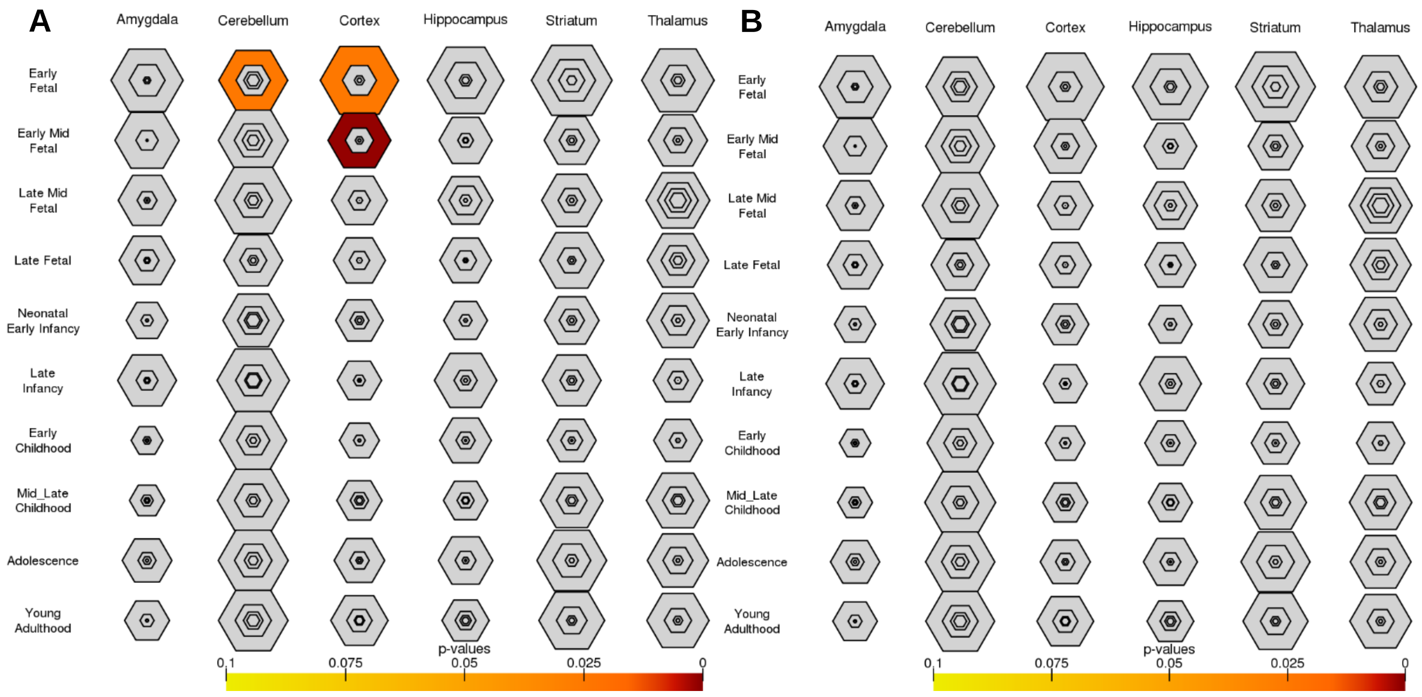


**Fig. S13: Cell-type-specific expression analysis.** A) 163 candidate genes with private LGD in probands only and B) 83 candidate genes with LGD in siblings only.

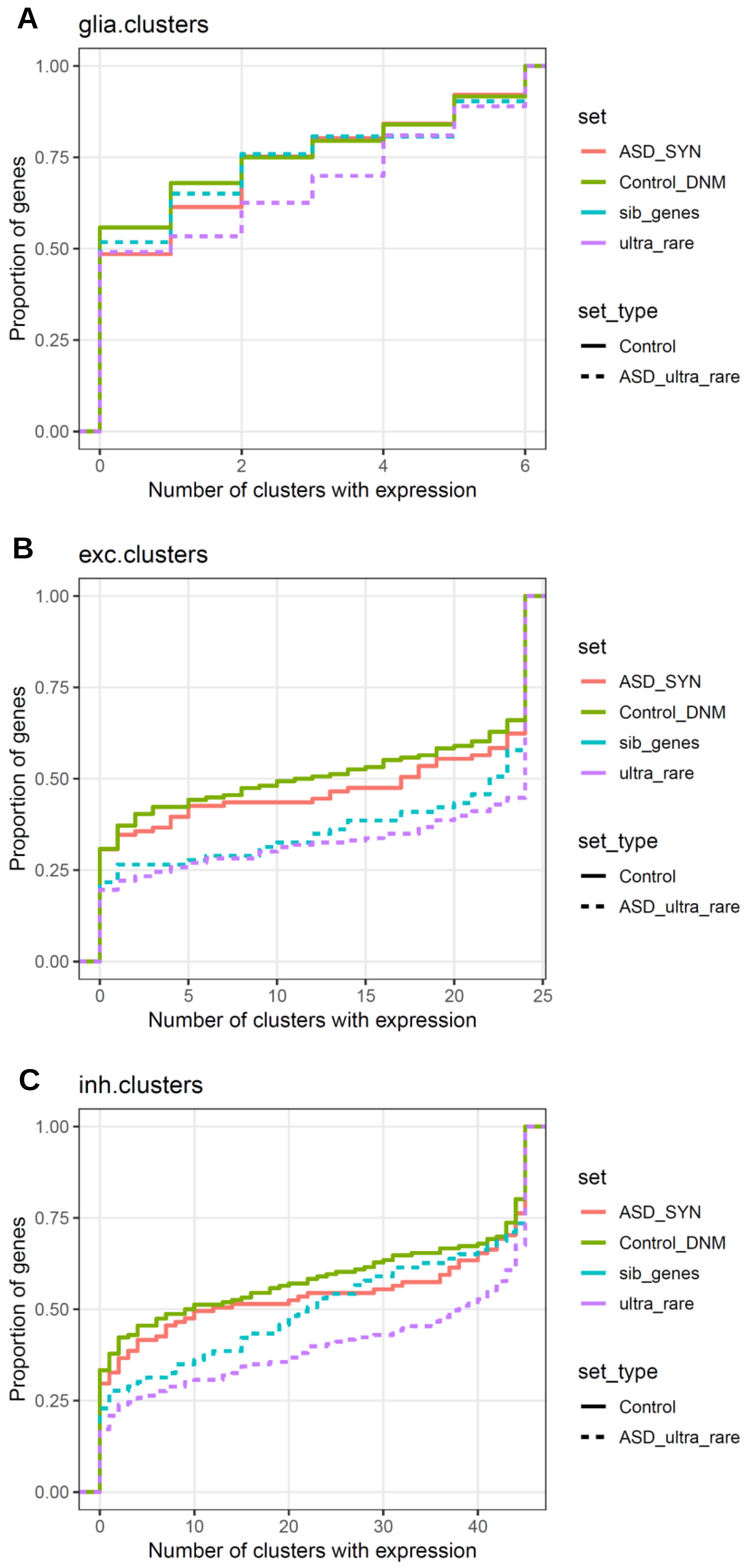


**Fig. S14: Proportion of candidate genes expressed brain cells.** A) Glia, B) excitatory, and C) inhibitory neurons. Excitatory and inhibitory neurons are enriched for expression of candidate genes as compared to control sets, but not in comparison to genes ascertained in siblings for the same criteria.

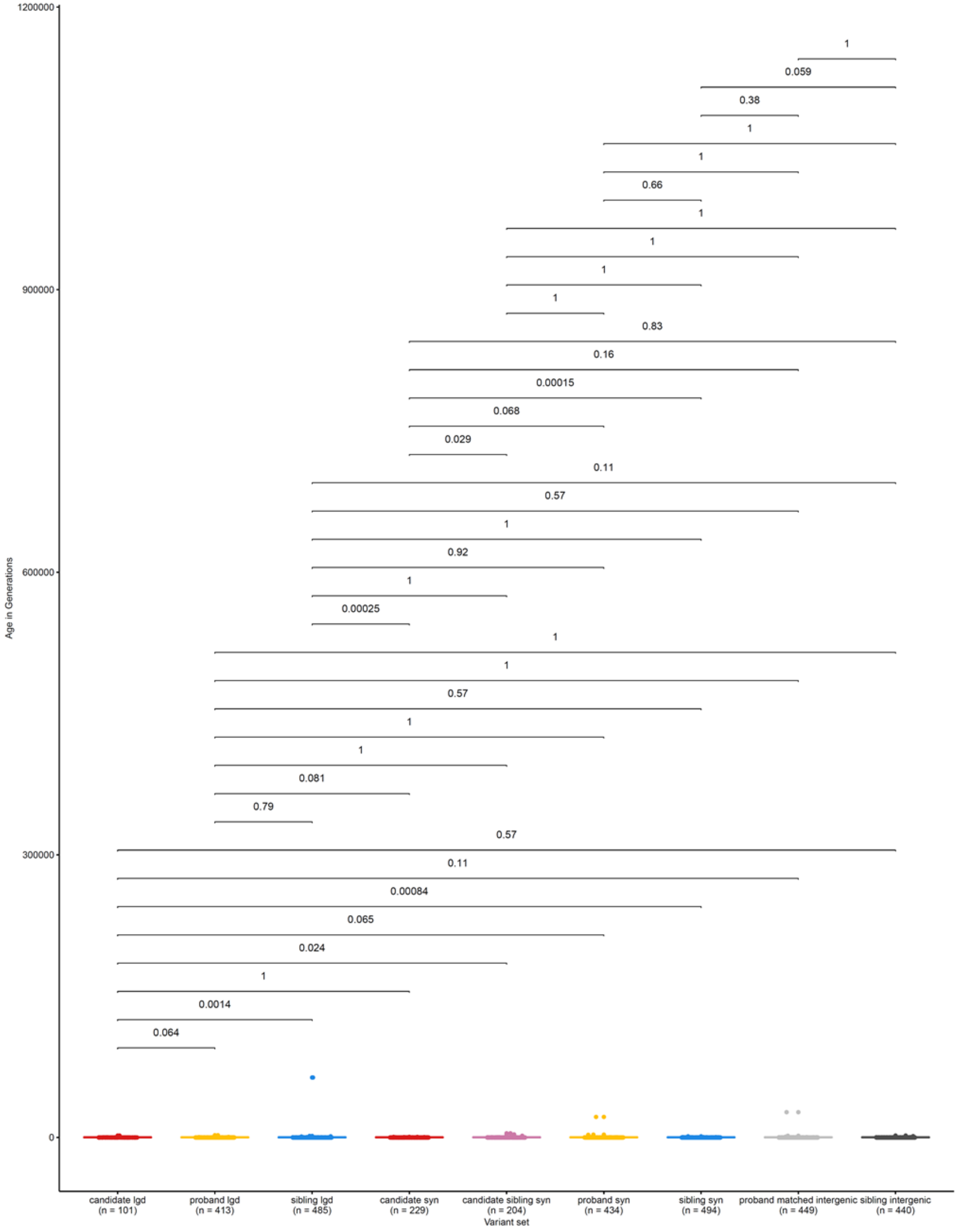


**Fig. S15: Allele age estimates and comparisons for LGD and SYN variants in the EUR subset of our discovery cohort.** P-values are Bonferroni-corrected for 36 tests.

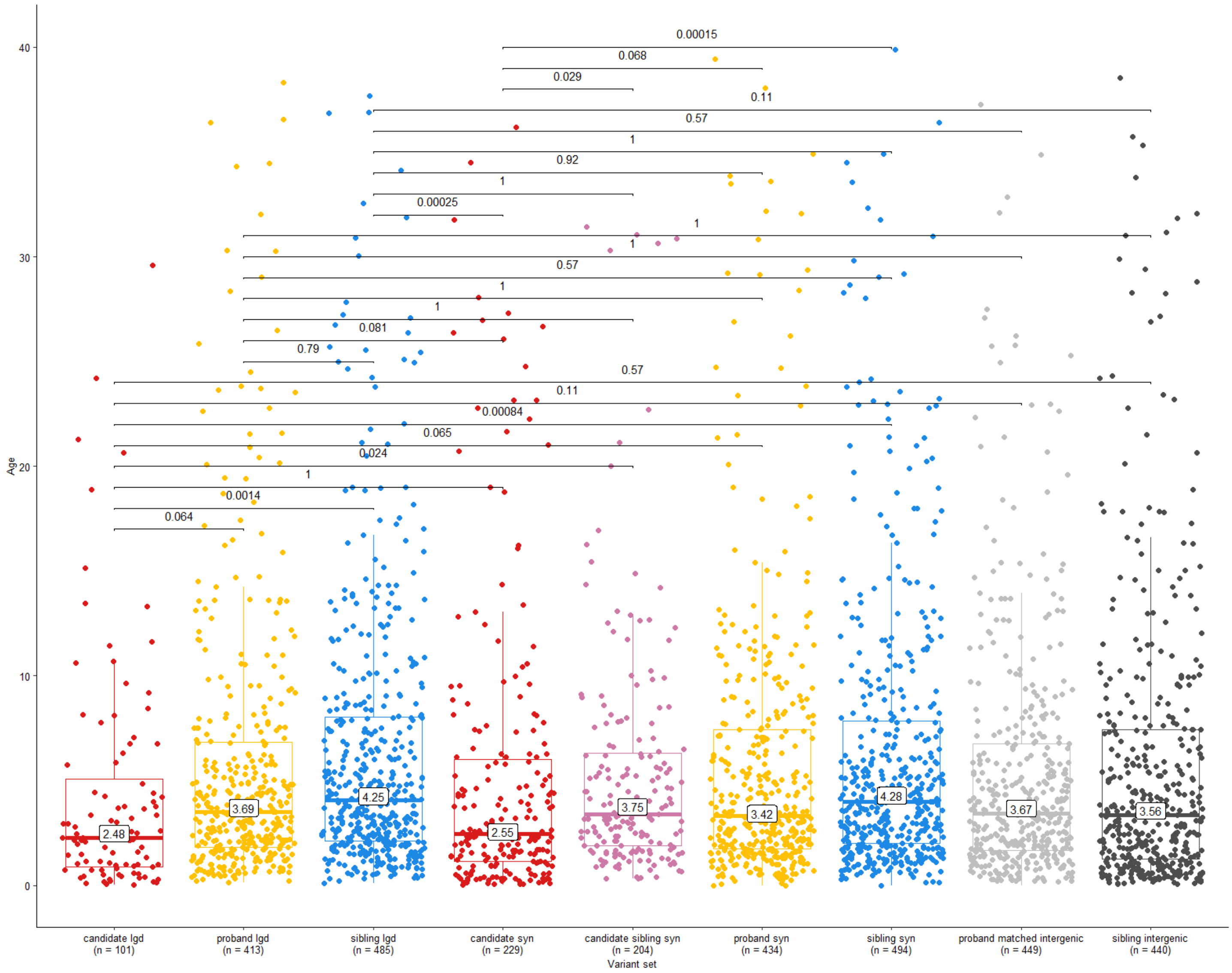


**Fig. S16: Allele age estimates for LGD and SYN variants within the EUR subset of our discovery cohort.** Plot is zoomed in to 20 generations (data older than this is included in calculating represented statistics (e.g., boxplots, medians, p-values) but is not visualized). Medians and p-values represent the full dataset and p-values are Bonferroni-corrected for 36 tests.

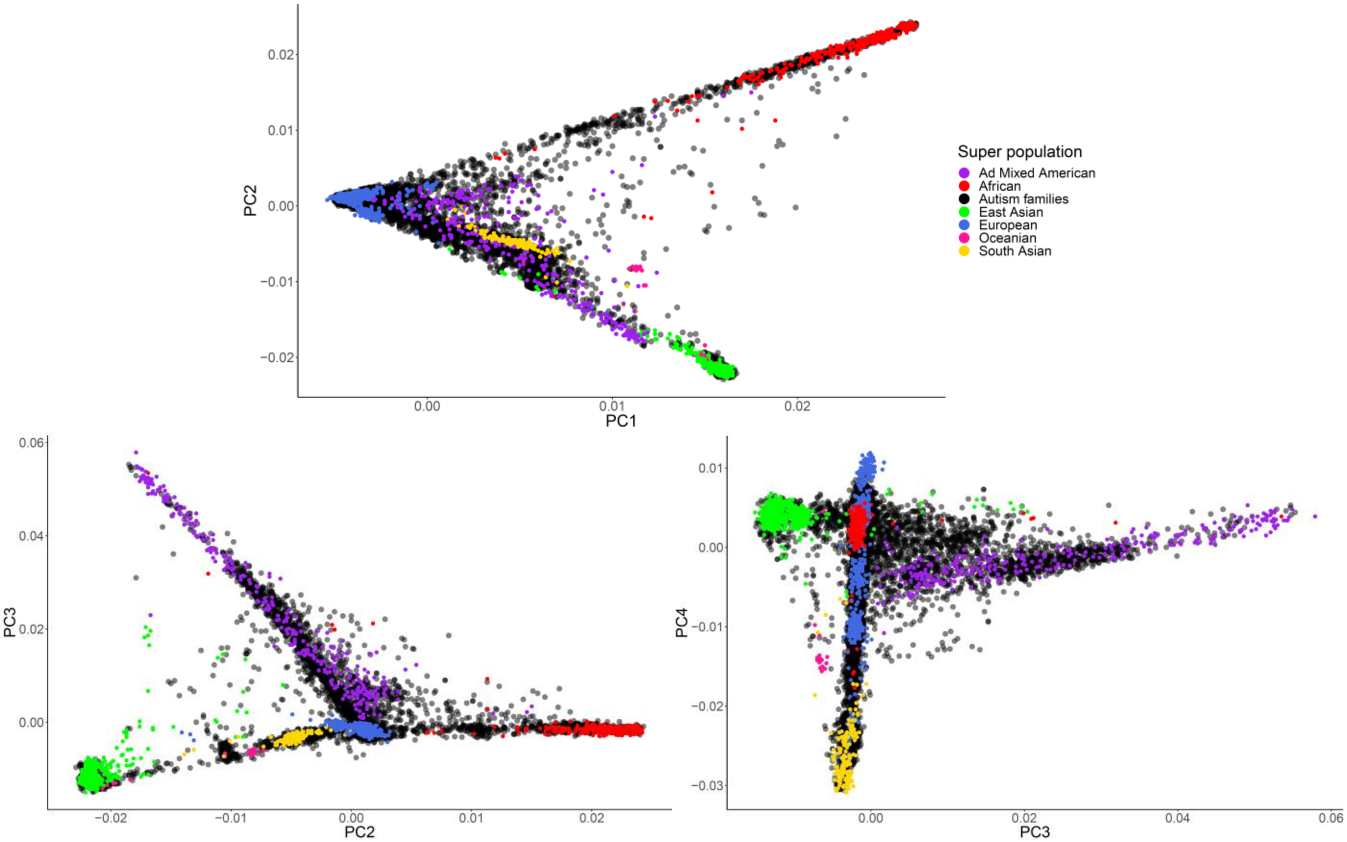


**Fig. S17: Plots of the first four principle components from PCA comprised of two reference cohorts, our discovery cohort, and our replication cohort.** Plots highlight the diversity of the cohorts.
